# TET CpG sequence context specific DNA demethylation shapes progression of IDH-mutant gliomas

**DOI:** 10.1101/2025.10.09.681314

**Authors:** Youri Hoogstrate, Santoesha A. Ghisai, Levi van Hijfte, Rania Head, Iris de Heer, Marta Padovan, Maurice de Wit, Wies R. Vallentgoed, Angelo Dipasquale, Maarten M.J. Wijnenga, Bas Weenink, Rosa Luning, Sybren L.N. Maas, Adela Brzobohata, Michael Weller, Tobias Weiss, Maximilian J. Mair, Anna S. Berghoff, Adelheid Wöhrer, Albert Jeltsch, Johan A.F. Koekkoek, Hans M. Hazelbag, Mathilde C.M. Kouwenhoven, Yongsoo Kim, Bart A. Westerman, Bauke Ylstra, Anneke M. Niers, Kevin C. Johnson, Frederick S. Varn, Roel G.W. Verhaak, Mustafa Khasraw, Martin J. van den Bent, Pieter Wesseling, Pim J. French

**Affiliations:** Department of Neurology, Erasmus MC Cancer Institute, Rotterdam, The Netherlands; Department of Neurosurgery, University Clinic Erlangen, Erlangen, Germany; Department of Oncology, Oncology 1, Veneto Institute of Oncology IOV-IRCCS, 35128 Padua, Italy; RCCS Humanitas Research Hospital, Via Alessandro Manzoni 56, Rozzano (Milan), Italy; Department of Pathology, Erasmus MC Cancer Institute, Rotterdam, The Netherlands; Department of Pathology, Leiden University Medical Center, Leiden, The Netherlands; Department of Neurology, Clinical Neuroscience Center, University Hospital Zurich and University of Zurich, Zurich, Switzerland; Division of Oncology, Department of Medicine I, Medical University of Vienna, Vienna, Austria; Department of Pathology, Neuropathology and Molecular Pathology, Medical University of Innsbruck, Innsbruck, Austria; Division of Neuropathology and Neurochemistry, Department of Neurology, Medical University of Vienna, Vienna, Austria; Institute of Biochemistry and Technical Biochemistry, Department of Biochemistry, University of Stuttgart, Stuttgart, Germany; Department of Neurology, Leiden University Medical Center, Leiden, The Netherlands; Dept. Of Pathology, Haaglanden MC, The Hague, The Netherlands; Department of Neurology, Amsterdam UMC, Amsterdam, The Netherlands; Department of Pathology, Amsterdam UMC, Cancer Center Amsterdam, Amsterdam, The Netherlands; Department of Human Genetics, Amsterdam UMC, Amsterdam, The Netherlands; Department of Neurosurgery, Yale University, USA; The Jackson Laboratory for Genomic Medicine, Farmington, CT, USA; Department of Genetics and Genome Sciences, University of Connecticut Health Center, Farmington, CT, USA; Institute for Systems Genomics, University of Connecticut, Storrs, CT, USA; Department of Neurosurgery, Duke University, Durham, North Carolina, USA; Princess Máxima Center for Pediatric Oncology, Utrecht, The Netherlands

## Abstract

**Background:** Treatment decisions in IDH-mutant oligodendrogliomas are shaped by tumor aggressiveness, underscoring the need for objective grading of these malignant brain tumors.

**Material and Methods:** We collected 302 primary and recurrent resections from oligodendrogliomas and performed Ki-67 staining, proteomics and DNA methylation profiling.

**Results & conclusion:** During tumor progression, DNA methylation of oligodendrogliomas changed along a continuum. This continuum is linked to increased epigenetic aging, methylation of transcription factors and Ki-67+ cell density, and to large scale DNA demethylation. Demethylation was correlated with CpGs flanking sequences preferred by TET enzymes. We confirmed these findings in previously profiled astrocytomas, indicating IDH-mutant gliomas progress along a shared epigenetic axis. We developed an objective DNA methylation based prognostic continuous grading coefficient (CGC^ψ^) that captured these changes and outperformed WHO grading for oligodendrogliomas. Our findings underscore the potential of DNA methylation-based grading to more accurately reflect tumor biology and inform clinical decision-making in IDH-mutant gliomas.

## Introduction

Oligodendrogliomas, IDH-mutant & 1p/19q-codeled (‘oligodendrogliomas’) are isocitrate dehydrogenase *IDH1*/*IDH2* mutated gliomas that are molecularly distinct from astrocytomas, IDH-mutant (‘astrocytomas’), based on the presence of a codeletion of chromosome arms 1p and 19q, often with mutations in the *TERT*-promotor and generally retained *ATRX* expression ^1,2^. With a median overall survival of 15 years, the group of patients diagnosed with oligodendroglioma has a more favorable outcome compared to those with other prevalent diffuse glioma types ^3,4^. Whereas tumor aggressiveness of oligodendroglioma is believed to be a continuum ^5^, in clinical practice a distinction is made between Central Nervous System World Health Organization (CNS WHO) grade 2 and grade 3. According to the WHO classification ^1^ this distinction is determined based on histological criteria such as presence/absence of high mitotic activity, microvascular proliferation, and necrosis. While this classification is used for treatment decision-making, the criteria for grading are not unequivocally defined ^6^, for instance by the lack of a standardized cut-off for cell division markers Ki-67/MiB or mitotic count ^5–8^. As a result, there is a high interobserver variability in oligodendroglioma grading ^9^. The difficulty in grading oligodendrogliomas is demonstrated by several recent studies in which no significant difference in prognosis between WHO grades was found ^10–12^, indicating a need for better and objective ways to define prognosis.

A limited number of imaging and (epi)genetic markers have been linked to the malignant progression of oligodendroglioma, but often not independently validated ^2,13–20^. These include contrast enhancement on MRI ^21,22^, the number of mitoses per mm^2 23,24^, HOX locus hyper-methylation ^25^, including *HOXD12* ^26^ and *HODX13* ^27^, and *CDKN2A/B* homozygous deletions ^18,28,29^. A subset of oligodendrogliomas, named “oligosarcoma”, develop an aggressive phenotype with sarcomatous features and are characterized by a distinct DNA methylation profile ^30^. Clinicians often face a dilemma: whether to defer radiotherapy and chemotherapy, or to pursue a more aggressive approach. These treatment decisions are based on the anticipated prognosis of patients, and therefore more objective robust stratification approaches are required to determine the aggressiveness of oligodendroglial tumors ^31–33^. Due to its widespread use in neuro-oncology, DNA methylation-based profiling has been proposed to enhance oligodendroglioma grading ^11,34^ and has been applied to assign prognostic features in gliomas ^25,30,34–39^.

Studies in which patients and their tumors are followed longitudinally can yield insight into the value of such markers, and into the molecular mechanisms underlying the malignant transformation of gliomas ^35,40–42^. We have therefore established the GLASS-OD workgroup as part of the International Glioma Longitudinal AnalySiS (GLASS) consortium ^43^. We collected longitudinal tumor samples from 127 patients diagnosed with oligodendroglioma that had undergone more than one surgical intervention with at least 6 months in between those interventions and investigated the molecular profiles of these tumors over time and grade.

## Results

### Primary – recurrent GLASS-OD oligodendroglioma cohort

For this study, DNA methylation data was generated from 267 surgical resections obtained from 127 patients. After removal of samples with low tumor purity (<10%) or poor quality, and patients with samples that lacked 1p/19q codeletions, the final discovery dataset consisted of 211 surgical interventions of 111 patients with oligodendroglioma from multiple institutions (**Fig. 1a-b, Table S1**). Clinical parameters including CNS WHO grade were provided by the respective hospitals. An independent oligodendroglioma validation set, comprising samples from patients who underwent single and multiple surgeries, was assembled from the literature and included 91 tumor samples from 76 patients ^11,30,44,45^ (**Supplementary Fig. 1a, Table S2**). From the TCGA-LGG dataset ^46^ 150 primary oligodendroglioma samples were obtained and from the GLASS-NL study ^35^ 203 primary – recurrent astrocytomas.

**Fig. 1.**
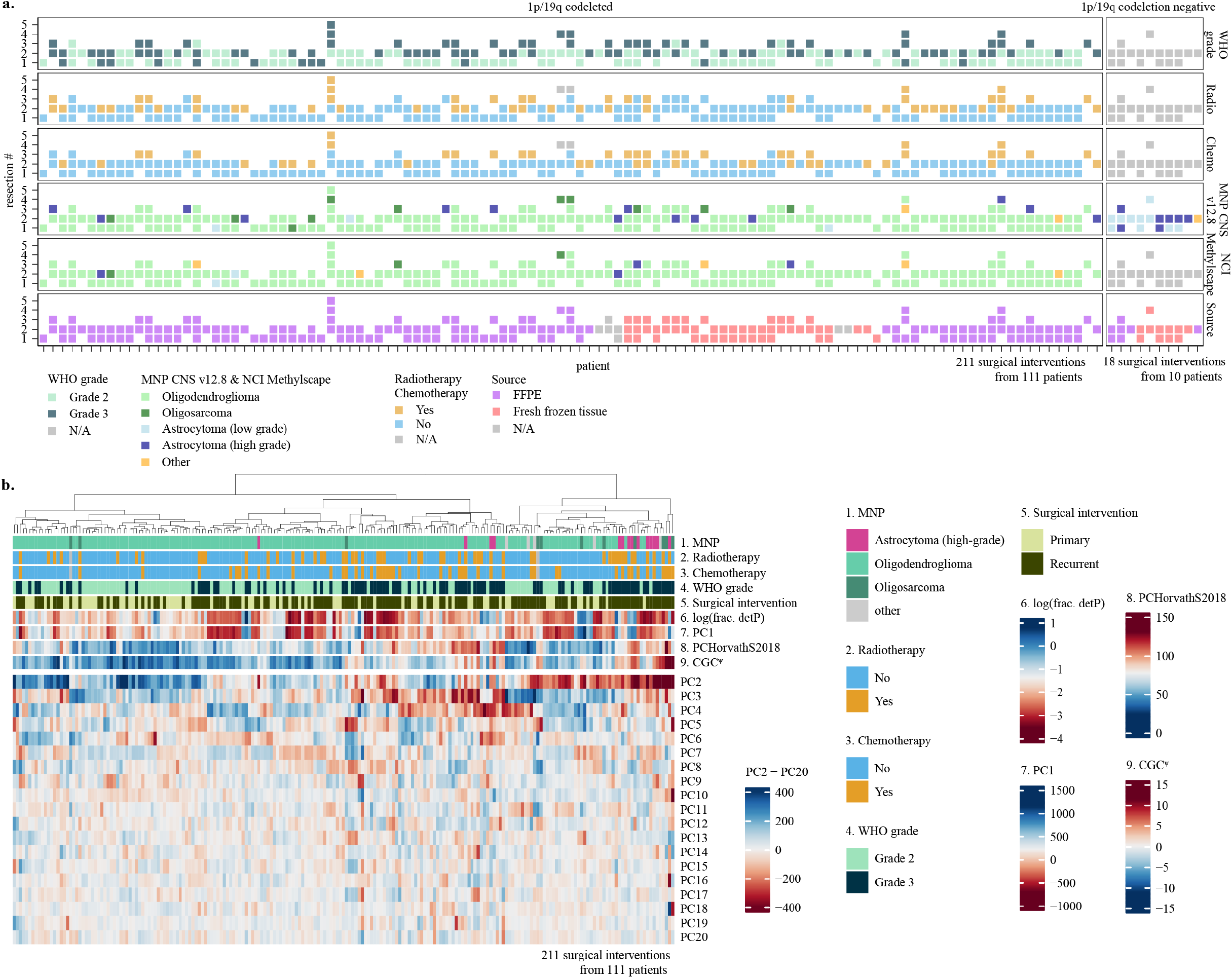
Cohort overview. **(a)** Overview of surgical interventions per patient in the GLASS-OD DNA methylation dataset. Patients were included based on histological diagnosis. Patients are separated in panels on their 1p/19q co-deletion status determined by DNA methylation data. The 1p/19q codeletion negative patients were excluded from further analysis. **(b)** Heatmap of n=211 1p/19q codeleted surgical interventions combined with patient and sample characteristics clustered on principal components PC2 - PC20.

### Oligodendrogliomas and astrocytomas progress along a shared epigenetic axis

To characterize oligodendroglioma methylomes along malignant transformation, we performed differential methylated probe (DMP) analyses. Aiming to better understand the evolutionary trajectories and the respective implications of CNS WHO grading, we compared both *primary* with *recurrent* tumors and *WHO grade 2* with *3* tumors. As there were patients with samples from more than 2 surgical interventions, we consistently compared primary tumor samples with the last available recurrence (**Supplementary Fig. 2a**). This maximizes the time between surgical resections (median: 67.3 months) and its effects, permitting resection comparisons of identical or even descending WHO grade. Comparing WHO grade 2 tumors with WHO grade 3 tumors maximizes the effect of malignant phenotypes as defined by neuropathological assessment. Likewise, in case patients had multiple surgical interventions of similar grade, the first grade 2 and/or last grade 3 was consistently chosen (**Supplementary Fig. 2a**). We compared changes between primary versus recurrent and found a large degree of DNA-demethylation at tumor recurrence. A similar, but larger demethylation was observed when comparing WHO grade 2 versus 3 (**Fig. 2a**).

**Fig. 2.**
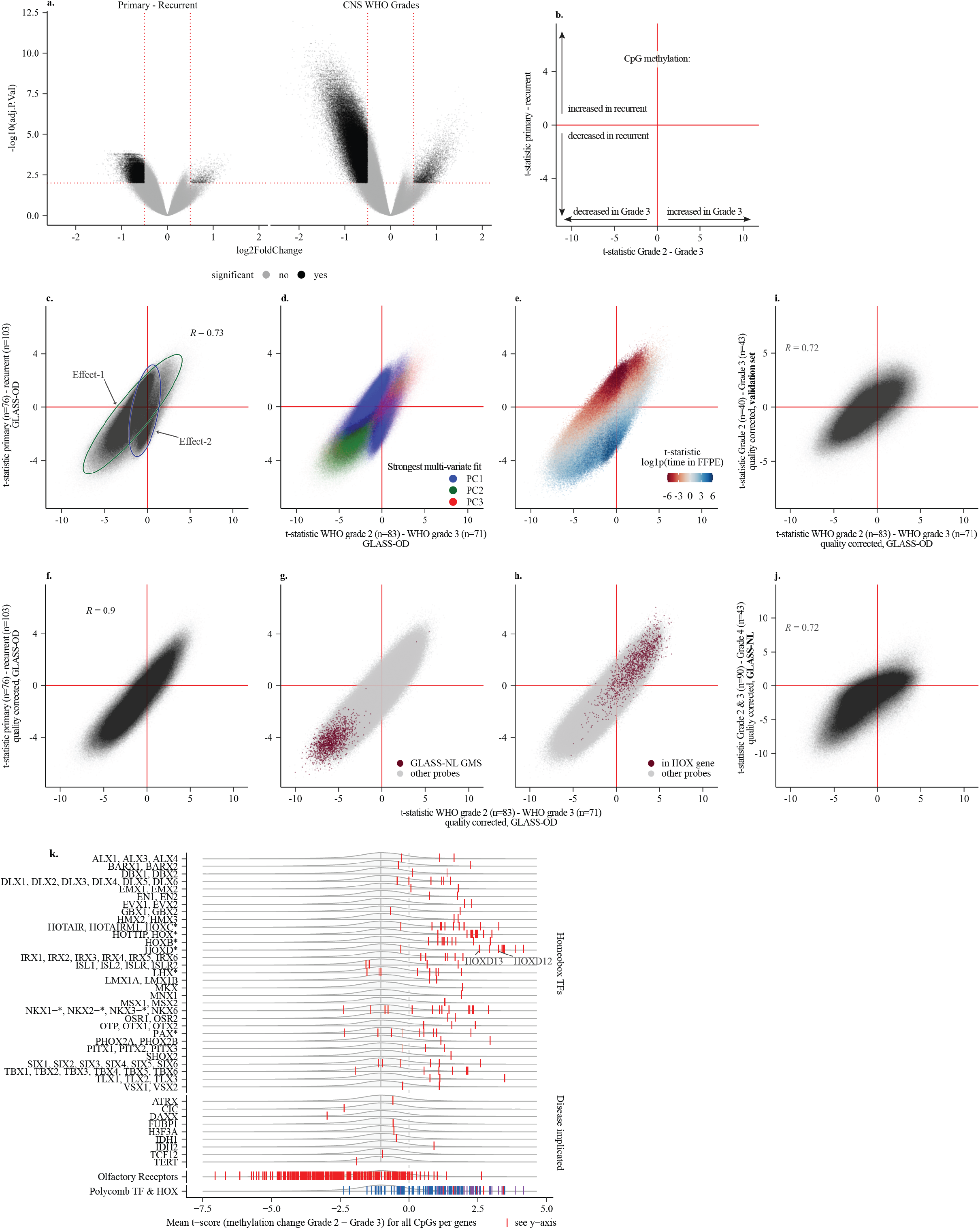
Multiple mechanisms contribute to DNA methylation differences between CNS WHO grades and between primary and recurrent tumors. **(a)** Volcano plots summarizing the DMP analyses, comparing the DNA methylation M-values per CpG between primary – recurrent tumor samples (left) and between CNS WHO grades (right). Significant CpGs (adj. P value < 0.01, |log2FC| > 0.5) are marked in black, non-significant CpGs are marked in gray. Red lines represent the significance thresholds. The y-axes, -log10(adjusted p-value), are scaled evenly. **(b)** The DMP analyses were integrated by their t-statistics, as t-statistics are signed like a logFC and normalized against standard error like a p-value. This panel provides a schematic representation for the interpretation of the axes for the upcoming panels. The x-axis represents the relative change in methylation observed between CNS WHO grades, where negative values indicate a decrease in methylation at grade 3 and positive values indicate an increase in methylation at grade 3. The y-axis represents the relative change in methylation observed between primary versus recurrent tumors, where negative values indicate a decrease in methylation in recurrent tumors and positive values indicate an increase in methylation in recurrent tumors. **(c)** Integration of the DMP analyses by their t-statistics comparing CNS WHO grades (x-axis) and primary–recurrent (y-axis) in the GLASS-OD dataset. Two optical effects are highlighted with ellipses (effect-1: green, effect-2: blue). **(d)** Same as (c), but colored by the principal component each CpG related strongest to in a multivariable model. **(e)** Same as (c), but colored by the t-statistic of an additional DMP model that related the CpG probes’ methylation to the duration each tissue sample was stored in FFPE. For fresh frozen samples the time was set to 0. **(f)** Integration of the quality adjusted DMP analyses by their t-statistics comparing CNS WHO grades (x-axis) and primary–recurrent (y-axis) in the GLASS-OD dataset. **(g)** Same as (f), but colored by the CpGs annotated within HOX genes. **(h)** Same as (f), but colored by the CpGs most differentially methylated between primary – recurrent astrocytomas from the GLASS-NL study. **(i)** Integration of the quality corrected DMP analyses comparing between WHO grades in the GLASS-OD dataset (x-axis) and validation set (y-axis). Pearson correlation coefficient of the t-statistics is indicated with R. **(j)** Integration of the quality corrected DMP analyses comparing between WHO grade 2 vs. grade 3 oligodendrogliomas (GLASS-OD) on the x-axis and WHO grades 2 & 3 vs. grade 4 astrocytomas (GLASS-NL) on the y-axis. Pearson correlation coefficient of the t-statistics is indicated with R. **(k)** Outcome of quality corrected DMP analysis comparing between WHO grade in oligodendrogliomas, aggregated at gene level. The x-axis represents the per gene aggregated t-statistics (x-axis), relative to the distribution of per-gene aggregated t-statistics for all genes (gray). The median of all genes is indicated with a gray vertical line. Individual homeobox TFs and other genes implicated in glioblastoma are indicated on the y-axis. At the bottom, genes from the olfactory receptor family and polycomb TFs and HOX genes are indicated.

We intersected the outcomes of both comparisons by their t-statistics, as these are signed like a LogFoldChange and normalized against standard error like a p-value (**Fig. 2b**). Their outcomes were correlated but also indicated two optical underlying differences (‘effects’) (*R*=0.73; **Fig. 2c**). To pinpoint individual factors underlying these effects, we performed principal component analysis (PCA). We mapped the strongest contribution of each CpGs to the first three components, showing that optical effect-2 was represented by CpGs that best fitted the first principal component (PC1) (**Fig. 2d**). For these CpGs, the difference in methylation was more pronounced in the primary versus recurrence comparison than between grades. It was associated with the per-sample fraction detection-P-value, a metric that represents probe data quality (**Supplementary Fig. 2b**). Given that effect-2 was more pronounced between primary and recurrent samples which encompassed the longest time intervals between resections compared to CNS WHO grades, we suspected that it represented a cytosine deamination artifact resulting from prolonged FFPE storage. To address this, we estimated each CpG’s association with the respective time the tissue was stored in FFPE (**Fig. 2e**). This displayed a similar overlap between optical effect2, PC1 and the association with detection-P. Effect-2 was characterized by an increase in methylation at recurrence of probes specifically matching the TA[CpG] sequence and loss of methylation at recurrence of probes with high CpG count, typically of probe type I (**Supplementary Fig. 2c-e**). These findings suggest that (methyl-)cytosine deamination is specific to the CpG’s surrounding sequence with apparent differences between deamination of CpGs and mCpGs. Since effect-2 is primarily driven by DNA quality, and therefore an undesired technical artifact rather than biologically relevant. We repeated the DMP analyses by incorporating quality associated PC1 to correct for this artifact. Incorporation of this factor almost entirely eliminated *effect-2* (**Fig. 2f**) and t-statistics between the intersected grade- and primary-recurrence comparisons were highly correlated (R=0.90). The CpG sites showing differential methylation were skewed towards genome-wide DNA methylation loss at tumor recurrence and grade 3 (p < 0.01, single-sided t-test comparing t-statistics with μ=0). This included the earlier reported CpGs with the largest change in DNA methylation between primary and recurrent astrocytomas (**Fig. 2g**) ^35^. Of the significant probes, 9.1% showed increased methylation at high grade (FDR corrected p-value < 0.01 & |log2FC| > 1), in which CpGs annotated within *HOX* genes were overrepresented (**Fig. 2h**). The overall differences were confirmed in an independent validation set (*R* = 0.72, p <0.01 chi-square test on significant probes, **Fig. 2i**). Large scale DNA-demethylation combined with increased methylation of *HOX*-gene CpGs has also been observed in IDH-mutant astrocytomas, where they also are associated with tumor grade and tumor recurrence ^25,35,47–49^. We therefore aimed to further investigate to what extent the malignant change in DNA methylation of oligodendrogliomas and astrocytomas show similarities. The comparison between grades in astrocytoma (CNS WHO grade 2 & 3 vs. grade 4, GLASS-NL dataset ^35^, both quality corrected) compared with oligodendrogliomas showed indeed that the changes in both tumor subtypes were correlated (*R* = 0.72, p <0.01 chisquare test on significant probes, **Fig. 2j**). These data indicate that both IDH-mutant glial tumor subtypes progress along a shared epigenetic axis.

DNA methylation changes in oligodendrogliomas between WHO grades were investigated at gene level. Virtually all the homeobox transcription factors (TFs) had increased mean methylation, as did polycomb associated TFs, including members of the reported *HOX* gene loci ^26,45,50^ (**Fig. 2k**). Conversely, genes of the olfactory receptor family were characterized by methylation loss. From genes implicated in oligodendroglioma, decreased methylation of *TERT* (q-value: 1.77e^-5^) and *ATRX* complex member *DAXX* (q-value: 2.32e^-19^) was observed as grade increased.

### WHO grade specific changes in methylation are TET sequence context-specific

Correlated outcomes within oligodendroglioma and between oligodendroglioma and astrocytoma indicate per-CpG specificity and suggest these changes in methylation are not a random process. To explore this further, we mapped the per-CpG change in methylation into bins of CpGs with identical surrounding flanking sequences (the sequence context). Interestingly, we find that the grade-associated DNA demethylation was sequence context specific (**Fig. 3a**). CpGs were more prone to DNA demethylation when flanked by 5’ AA or 3’ TT sequences and more stable in the context of multiple CG di-nucleotides. Enzymes from the DNA Methyltransferase (DNMT) and Ten-eleven translocation methylcytosine dioxygenase (TET) families maintain DNA methylation and exhibit preferences for flanking sequences ^51,52^. We hypothesized that if such a mechanism is altered, this would affect DNA methylation in a sequence context specific manner. To address this, we correlated the per sequence context methylation change between CNS WHO grades with the flanking sequence dependent activities of DNA (de)methylating TET1-3 and DNMT1-3 enzymes ^51–58^ (**Fig. 3b-h**). This revealed a strong correlation between demethylation patterns and TET DNA-demethylation flanking sequence preferences (R = –0.73 ∼ –0.83). After applying a correction to the quality effect specific to TA[CpG]NN contexts (**Supplementary Fig. 3a**), the sequence context-specific demethylation in the astrocytoma dataset displayed a similar correlation with TET enzyme flanking sequence preferences (R = –0.71 ∼ –0.86, **Supplementary Fig. 3b**). These findings demonstrate that grade-associated DNA-demethylation in both oligodendrogliomas and astrocytomas are specifically stronger at sites preferential to TET DNA-demethylating enzyme activity.

**Fig. 3.**
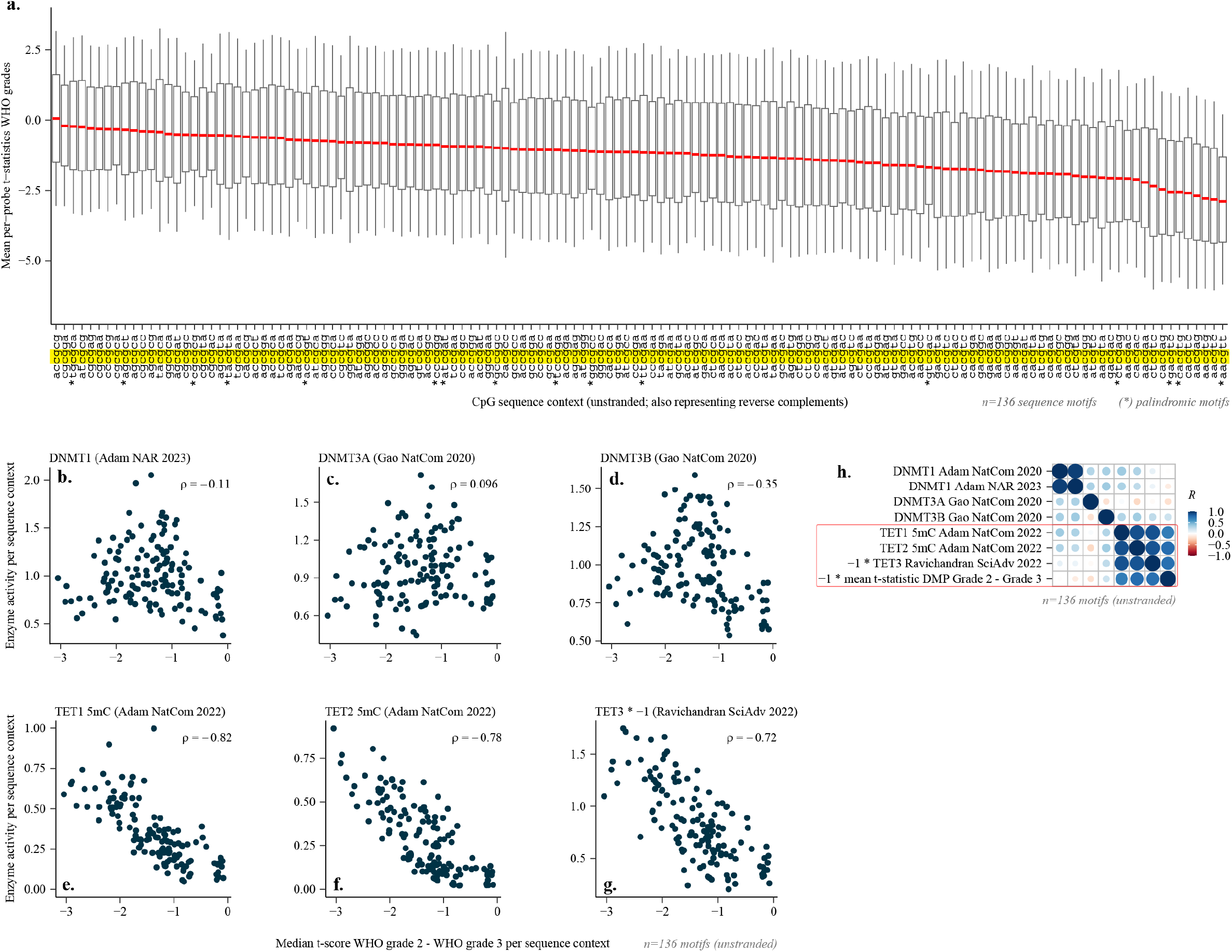
Demethylation is strongest in CpGs with flanking sequences preferred by TET. **(a)** The mean t-statistic (between WHO grades, GLASS-OD, quality-corrected; y-axis) of all CpGs grouped per sequence context (x-axis). Sequence contexts are aggregated in an unstranded manner, thus also representing their reverse complements. Contexts with identical reverse complements are indicated with an asterisk (*). **(b, c, d, e, f, g)** Spearman correlation coefficients (ρ) and scatterplots for the mean t-statistics per sequence context compared with the (de)methylation preferences of TET and DNMT enzymes. These preferences were obtained from the literature. **(h)** Pearson correlation plot with all per-sequence context metrics from above combined.

### CGC^ψ^: molecular continuous grading of IDH-mutant oligodendrogliomas

To capture malignancy of IDH-mutant astrocytomas reflecting its continuous nature ^35,59^, we previously developed a DNA methylation based Continuous Grading Coefficient (CGC) ^25^. As the underlying probabilities used by CGC are entangled with tumor subtype classification, CGC is astrocytoma specific and does not generalize to other tumor subtypes. Because oligodendrogliomas showed shared temporal changes in methylation compared to astrocytomas, we wanted to assess the presence and prognostic value of CGC in oligodendrogliomas. To this end, we aimed to define a predictor of this grading continuum that is independent of tumor subtype classification. We trained a Least Absolute Shrinkage and Selection Operator (LASSO) regression model on the GLASS-NL IDH-mutant astrocytoma samples, using methylation M-values directly to predict the calculated CGC coefficient for these samples. We used 10-fold cross-validation to assess the performance of predicting CGC, achieving a Relative Root Mean Squared Error (RRMSE) of 0.352 and Pearson correlation of *R*=0.94 (**Supplementary Fig. 4a**). The final model trained on all GLASSNL astrocytoma samples (CGC^ψ^) consisted of n=168 predicting CpG probes. Among the genes annotated to these CpGs were *WNT1, MAPK3, HOXA3, HOXA6, HOXA7, HOXA9*, and *HOXC12* (**Table S3**).

We applied CGC^ψ^ to GLASS-OD, and found the range of scores in oligodendroglioma to be higher than those in IDH-mutant astrocytoma (p=0.015, Wilcoxon test, **Supplementary Fig. 4b**). To get an indication of how CGC^ψ^ relates to the differences between primary–recurrent tumors and WHO grade in oligodendroglioma, we first estimated the per-CpG association with CGC^ψ^. We then color-coded the integrated DMP outcome accordingly and observed that the CpGs with the largest differences exhibited the strongest association with CGC^ψ^ (**Fig. 4a**). In GLASS-OD, CGC^ψ^ was significantly higher in recurrent tumors, CNS WHO grade 3, MNP CNS and NCI methylscape high-grade classes, and in the presence of CDKN2A/B homozygous deletions (p<0.01 [1.52e^-3^– 1.4e^-11^]), but did not differ between FFPE and fresh-frozen samples (p=0.38, **Fig. 4b**). In the validation set, CGC^ψ^ was also significantly higher in CNS WHO grade 3 tumors (p=6.01e^-5^, **Fig. 4c**). The mean CGC^ψ^ further increased with successive surgical interventions (**Fig. 4d**). These results confirm shared mechanisms of malignant progression in both IDH-mutant tumor types, following a continuum.

**Fig. 4.**
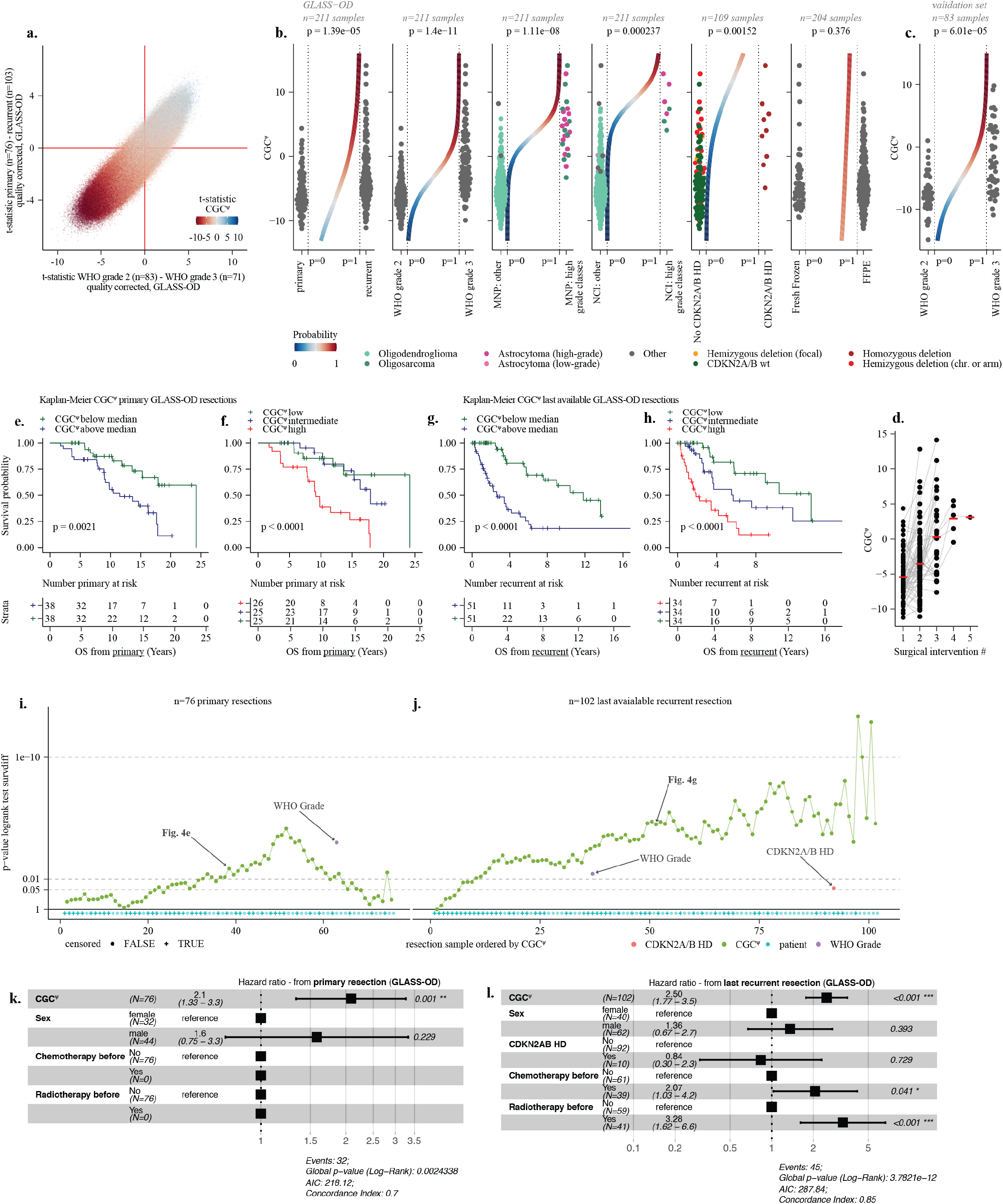
CGC^ψ^ is an objective prognostic continuous grading coefficient for oligodendrogliomas. **(a)** Integrated DMP plot displaying the relation of CGC^ψ^ with the changes in methylation between WHO grades and primary–recurrent. It is the same as (Fig. 2c), but colored by the t-statistic of an additional DMP model that related the CpG probes’ methylation to CGC^ψ^. **(b)** Scatterplots displaying the CGC^ψ^ across various conditions, with a logistic curve in the center. The p-values represent the logistic curve. Conditions from left to right: primary–recurrent, CNS WHO grades, MNP CNS classifier: astrocytoma (high grade) & oligosarcoma vs. other, NCI Methylscape classifier: astrocytoma (high grade) & oligosarcoma vs. other, *CDKN2A/B* status at last available resection, storage conditions (fresh frozen DNA or FFPE). **(c)** Same as (b) for WHO grade in the validation set. **(d)** Scatterplot displaying the temporal mean CGC^ψ^ per subsequent surgical intervention. **(e-h)** Kaplan-Meier estimates of the overall survival (OS) in the GLASS-OD dataset, splitting either the primary or last available recurrent tumor samples by the mean CGC^ψ^. In either two or three groups. **(i**,**j)** Cut-off sweep of p-values from Kaplan-Meier estimates of overall survival (OS) in the GLASS-OD dataset, ranked by CGC^ψ^. Primary tumors (i) and last available recurrent resections (j) were separated. For each cut-off along the CGC^ψ^, a log-rank test was performed. P-values for WHO grade and *CDKN2A/B* status (recurrent only) are indicated for reference. Patient censoring is shown at the bottom of each panel. **(k**,**l)** Forest plots of multivariate Cox proportional hazards (CoxPH) models for (k) overall survival from primary resection and (l) post-recurrence survival from the last recurrent resections in the GLASS-OD dataset.

Splitting the GLASS-OD cohort by their median CGC^ψ^ for either the primary or the last available resections resulted in CGC^ψ^-high groups with significantly worse survival (p=0.0021; log-rank test; **Fig. 4e,f**). When splitting into three equally sized groups by ranked CGC^ψ^ score, it displayed a distinct prognosis at recurrence, while in samples from the primary surgery CGC^ψ^ high had a significantly worse outcome from intermediate and low (**Fig. 4g,h**).

To test the prognostic value for each possible cutoff combination, samples were ranked by CGC^ψ^ and the overall survival difference for each cutoff was tested (**Fig. 4i,j**). Cox Proportional Hazard (CoxPH) regression confirmed that in both primary (p < 0.001; HR=2.1, 95% CI: 1.33 – 3.3; **Fig. 4k**) and recurrent tumors (p < 0.001; HR=2.5, 95% CI: 1.77 – 3.5; **Fig. 4l**) CGC^ψ^ was a significant predictor of poor prognosis. Prognosis of CNS WHO grade was however significant for primary (p<0.001; HR=3.8, CI=1.75 – 8.4, **Supplementary Fig. 4d**) but not for recurrent tumor samples (p=0.17; **Supplementary Fig. 4f**). In primary tumors, the concordance index of the model that included CGC^ψ^ (0.7, **Fig. 4k**) was only marginally higher than CNS WHO grade (0.69, **Supplementary Fig. 4d**). In primary tumors, CGC^ψ^ and WHO grade lost significance when included together, indicating they competed and had a similar performance (**Supplementary Fig. 4c**). In the recurrent tumors, the prognostic value of CGC^ψ^ (p < 0.001; HR: 2.5, 95% CI: 1.77 – 3.5; p-value; **Fig. 4l**) was superior to WHO grade (p = 0.17, **Supplementary Fig. 4f**), and performed independent of WHO grade (p < 0.001; **Supplementary Fig. 4e**). The models including CGC^ψ^ had a higher concordance index (0.85) than WHO grade alone (0.79), showing CGC^ψ^ had superior performance. CoxPH was performed on the last available samples of recurrent tumors in the validation set. Despite the limited number of samples with survival data, CGC^ψ^ was associated with survival (p=0.017, concordance index: 0.67, **Supplementary Fig. 4g**) while WHO grade was not (p=0.327; concordance index: 0.58; **Supplementary Fig. 4h**). To further validate CGC^ψ^ in primary tumors, we created a CGC^ψ^-derived model trained only on the intersection of probes present on both the 450k and 850k arrays. Applying this CGC^ψ/450k^ model to the primary oligodendroglioma samples of the TCGA-LGG dataset demonstrated more prognostic value (p<0.001, CoxPH, concordance index: 0.77, **Supplementary Fig. 4i**) than WHO grade (p=0.002, CoxPH, concordance index: 0.63; **Supplementary Fig. 4j**).

Oligodendroglioma samples with sufficient tissue available were stained for Ki-67 (n=111 with matching array). Cells were computationally detected and classified for Ki-67 positivity, resulting in positive cell fractions from <1.0% up to 48.0%, with 67.6% of the samples below 5% positive (**Fig. 5a,b,c,d,e**). Both the Ki-67 positive cell fraction and the density of positive cells per cm^2^ were significantly higher for both samples from CNS WHO grade 3 and recurrent tumors (p<0.0027; **Fig. 5f,g**). The number of Ki-67 positive cells per cm^2^ was a significant predictor of post recurrent survival (p=0.004; concordance index = 0.75; **Fig. 5h**). After incorporation of CGC^ψ^ into the model, Ki-67 lost significance and CGC^ψ^ outperformed Ki-67 (p<0.001; concordance index = 0.83; **Fig. 5i**).

**Fig. 5.**
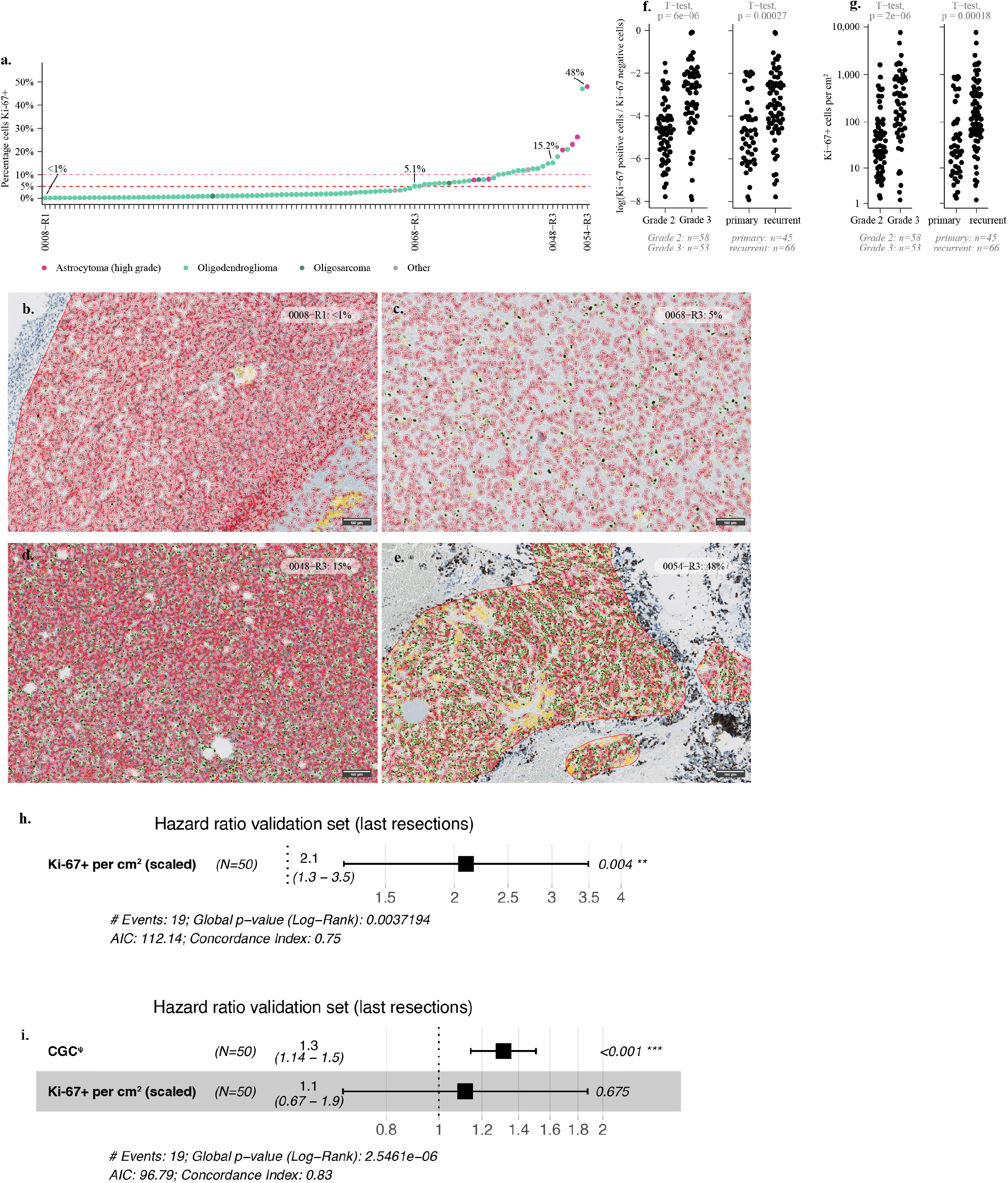
Ki-67 positive cell density is a prognostic factor in oligodendroglioma. **(a)** Percentage of Ki-67 positive cells relative to negative cells, determined by computational analysis of n=111 stainings. Tumor samples are shown on the x-axis, and the percentage of Ki-67 positive cells on the y-axis. **(b–e)** Representative screenshots of computational cell detection and classification of Ki-67 positivity. The percentage of positive cells and sample identifiers are indicated in the top right corner. Detected cells classified as Ki-67 positive are marked in green, Ki-67 negative in red, and artifacts in yellow. **(f)** Comparison of the log-transformed Ki-67 positive cell ratio across WHO grades (left panel) and between primary and recurrent tumors (right panel). P-values from two-sided t-tests are shown at the top. **(g)** Comparison of Ki-67 positive cell density (cells/cm^2^) across WHO grades (left panel) and between primary and recurrent tumors (right panel). P-values from two-sided t-tests are shown at the top. **(h)** Univariate Cox proportional hazards model on overall survival (CoxPH) assessing the prognostic value of Ki-67 density at the time of the last available surgical intervention.

### Accelerated epigenetic aging contributes to the axis of progression

Cellular aging and the cell replicative history within living tissue result in distinct DNA methylation patterns. There are several algorithms able to predict age and cell replicative history (epigenetic clocks) by making use of these patterns ^60^. We applied algorithms provided by metapackage dnaMethyAge ^61^ and clustered their outcome with other parameters (**Fig. 6a**, left panel). This was extended with differential comparisons between WHO grade and primary – recurrent (**Fig. 6a**, center and right panel). Of the three resulting clusters, virtually all clocks cluster together. All but two clocks were increased in WHO grade 3 (largest difference in *PCHorvathS2018*, linear regression: q=7.06e^-7^). Whereas it is self-evident that recurrent tumors are epigenetically older than their primary, closer inspection of *HorvathS2018* revealed predicted epigenetic ages at recurrence in some cases exceeding 100 years (**Fig. 6b,c**), older than their actual age. We observed that epigenetic age indeed exceeded patients’ chronological age at surgical intervention and accelerated specifically as CGC^ψ^ increased (**Fig. 6d**).

**Fig. 6.**
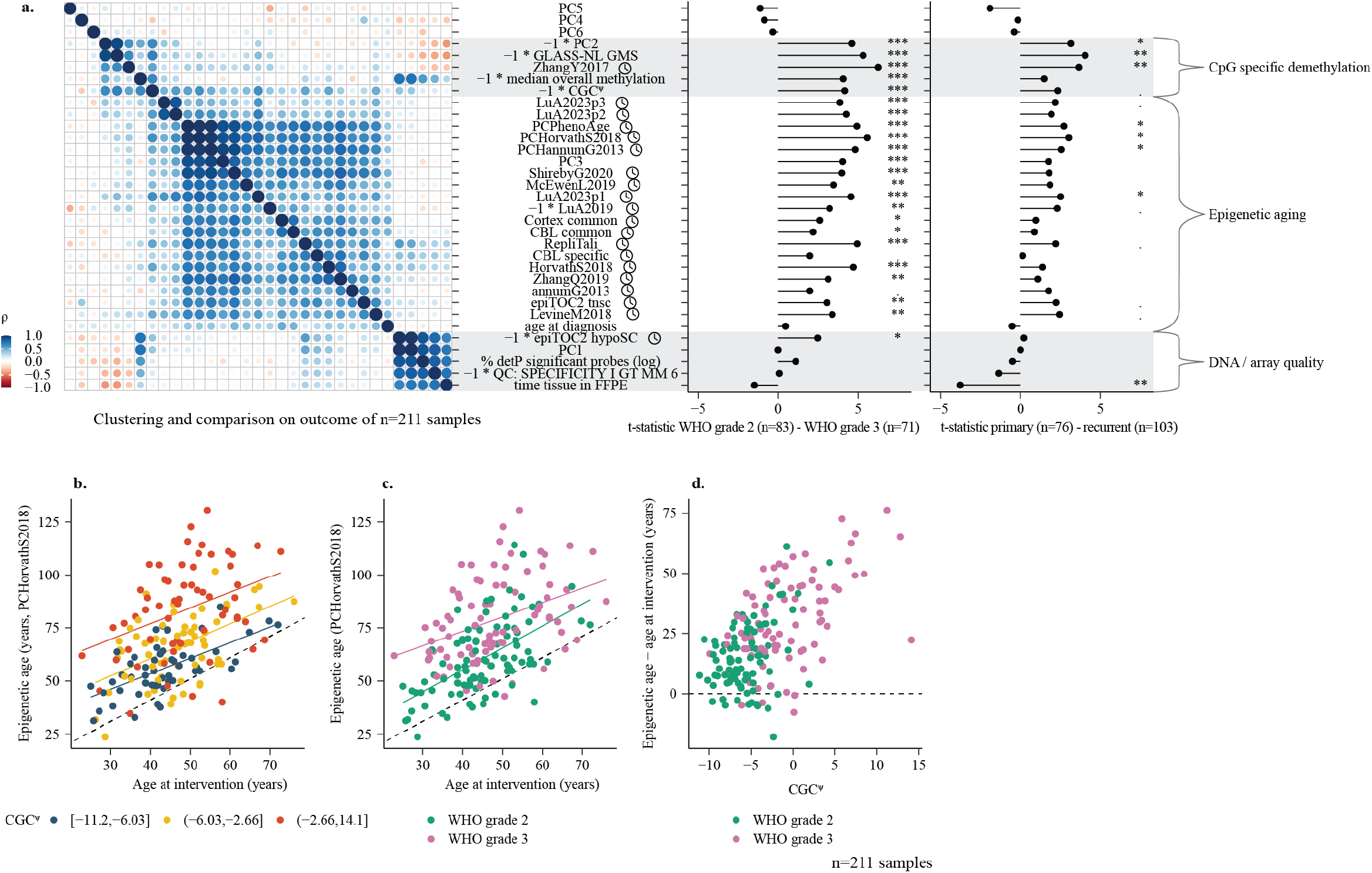
Epigenetic aging is increased as CGC^ψ^ increases. **(a)** Left: Correlation-based clustering of epigenetic clock outputs combined with other sample-level parameters in the GLASS-OD dataset. Right: Linear regression analysis comparing epigenetic clock estimates and additional sample metrics between primary – recurrent and WHO grades. The x-axis displays t-statistics from linear models comparing metrics across WHO grades (center) and between primary and recurrent tumors (right). False discovery rate (FDR)-corrected p-values (q-values) are indicated by asterisks (., *, **, *** for increasing significance). Spearman correlation coefficients (ρ) are shown where applicable. **(b)** Concordance between the chronological age at surgical intervention (x-axis) and the epigenetic age (PCHorvathS2018) derived from DNA methylation profiles (y-axis). Regression lines are shown for three CGC^ψ^ groups defined using the cut function in R. **(c)** Same as (b), but regression lines and colors by WHO grade. **(d)** Scatterplot of the actual difference between chronological and epigenetic age (PCHorvathS2018, y-axis) across CGC^ψ^ values (x-axis).

Of the principal components, PC2 and PC3 both differed significantly between CNS WHO grade and primary – recurrent and both correlated with CGC^ψ^ (**Fig. 6a, Fig. 2d**). This indicated that beyond quality, two underlying factors contribute to the changes in methylation, both captured by CGC^ψ^. PC3 clustered between the epigenetic clocks and displayed strong correlation with *PCHorvathS2018* (ρ=0.87, Spearman correlation). We further investigated the contribution of all polycomb TF-associated probes to PC3 (**Supplementary Fig. 5a,b**), and found CpGs within these genes contributed to PC3. For the subset of samples with matching Ki-67 stainings, we estimated correlation with Ki-67 density and positivity fractions with PC2, PC3 and the Horvath epigenetic clock. This revealed that Ki-67 positive cell density was in particular correlated with PC3 and the Horvath epigenetic clock (ρ=0.61–0.68; **Supplementary Fig. 5c**). Of the two underlying mechanisms, the changes represented by PC3 are related to accelerated epigenetic aging and linked with cell cycling, marked by increased methylation of a broad set of transcription factors, including those from the HOX loci, whereas PC2 is related to CpG demethylation.

### Oligodendroglioma progression is characterized by chromosomal losses

Associations were estimated between per-bin copy-number variations (CNV) with primary – recurrent, CNS WHO grades, CGC^ψ^, principal components 2 and 3 and post recurrence survival. Losses at chromosomes 4q, 9p, 13, 14q, 15 and 18 were associated with WHO grade (**Supplementary Fig. 6a**). When fitting to CGC^ψ^ instead of WHO grade, similar but more pronounced associations were detected (**Supplementary Fig. 6b**). Interestingly, when fitting to PC2 and PC3, certain genomic alterations fitted uniquely to either PC2 or PC3 (**Supplementary Fig. 6c,d**), suggesting these events contribute to distinct molecular mechanisms. Genomic losses of chr11p showed a trend towards post recurrence survival (**Supplementary Fig. 6e**).

We observed a homozygous loss of the *CDKN2A/B* locus in 9.9% (10/111) at the last available tumor sample. Homozygous *CDKN2A/B* deletions were typically focal and spanning the locus specifically. Hemizygous *CDKN2A/B* deletions were more frequent (35/111; 31.5%) and typically encompassed large genomic regions (**Supplementary Fig. 7a**) as they were mostly partial arm, whole arm or entire chr9 losses (**Table S4**). The 9p arm losses were found in 26/111 patients (25.7%) of which four also had a homozygous deletion event. Although the incidence was low (n=10), in multivariable analysis on the last available recurrent tumor (n=101), *CDKN2A/B* HD did not show a statistically significant different overall survival (**Fig. 4l**).

### Oligosarcomas are an aggressive subtype of oligodendroglioma with a lower tumor cell fraction

We uploaded our DNA methylation data of the GLASS-OD cohort onto the (Molecular NeuroPathology; MNP CNS; v12.8) methylation-based tumor classifier v12.8 ^36^. According to this classifier, 14 oligodendroglioma recurrences were high-grade astrocytoma (A_IDH_HG, **Fig. 1a**), of which six with a prediction confidence >= 0.84. As indicated in the cohort description, these samples had a 1p/19q codeletion (**Supplementary Fig. 8a**) and the classifier labeled tumors in the prior resections of these patients as oligodendroglioma. Furthermore, of the 11 samples the classifier v12.8 labeled as oligosarcoma, 7 were classified as high-grade astrocytoma in v11.b4, the last version before oligosarcoma was incorporated (v11.b4; **Supplementary Fig. 8b**). Oligodendrogliomas classified as either oligosarcoma or highgrade astrocytoma were unanimously characterized by the highest CGC^ψ^ scores (p=1.11e^-8^, t-test, **Fig. 4b**). NCI MethylScape classified four recurrent tumors to be high grade astrocytoma and three oligosarcoma, which were also characterized by high CGC^ψ^ (p=2.3e^-4^, **Fig. 4b**).

Our observation that oligodendrogliomas classified as high grade astrocytoma were characterized by a high CGC^ψ^ could indicate that aggressive astrocytomas and oligodendrogliomas converge towards an indistinguishable overarching epigenetic state. To test this, Uniform Manifold Approximation and Projection (UMAP) was performed and showed that oligodendroglioma methylation profiles remain distinguishable from astrocytoma (**Supplementary Fig. 8c**). Given that oligodendrogliomas remain distinct from astrocytomas, the observation that the methylation-based classification of oligodendrogliomas with high CGC^ψ^ branches into either high-grade astrocytoma or oligosarcoma, could indicate diverging evolutionary paths. However, when comparing methylation profiles between the two groups, no CpGs were differentially methylated after multiple-testing correction (**Supplementary Fig. 8d**). We did find that oligodendrogliomas classified as high grade astrocytoma were characterized by a significantly higher tumor cell fraction than those classified as oligosarcoma (p=0.009, t-test, **Fig. 7a,b**). To further investigate whether this branched classification fate could be attributed to tumor purity, we performed an *in silico* dilution experiment. Using our software package *idat-tools* (https://github.com/yhoogstrate/idat-tools), we simulated decreasing tumor purity by incrementally spiking in methylation data from non-tumor from whole-brain tissue ^62^ samples. In all four tested high-grade astrocytoma classified oligodendroglioma samples (0017-R3, 0008-R2, 0121-R3 & 0054-R3), incremental fractions of methylation data indeed changed the classification fate towards an oligosarcoma diagnosis, until it reached a non-tumor classification (**Fig. 7c, Table S5**). This change in classification was despite the spiked-in non-tumor samples do not fully recapitulate tumor micro-environments. These results indicate that tumor purity plays a role in classifying aggressive oligodendrogliomas as high-grade astrocytomas.

**Fig. 7.**
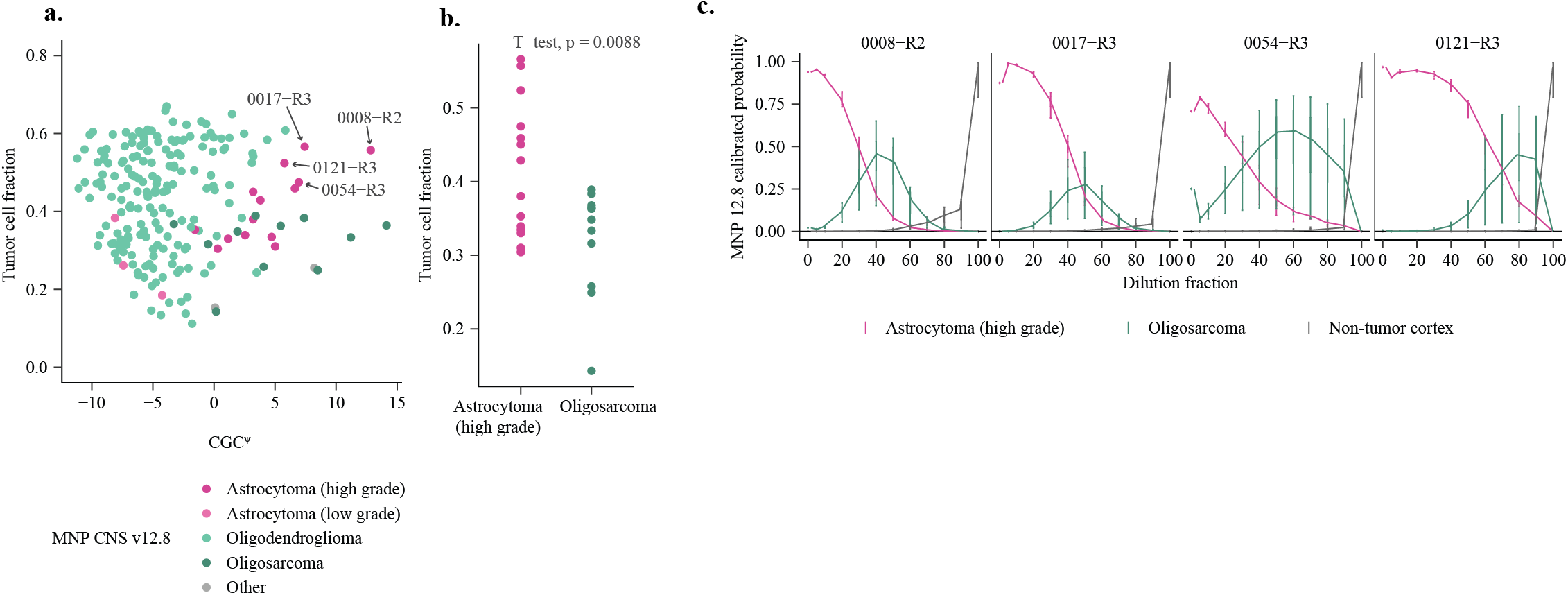
Classification of CGC^ψ^ high tumor as oligosarcoma or high-grade astrocytoma is related to tumor purity. **(a)** Scatterplot of GLASS-OD samples showing predicted CGC^ψ^ values (x-axis) versus tumor purity (y-axis). Samples are color-coded by MNP CNS predicted class. Samples selected for further analysis are highlighted. **(b)** Tumor purity comparison between GLASS-OD samples classified as oligosarcoma or high-grade astrocytoma. The y-axis represents tumor purity, and the p-value from a t-test is on top. **(c)** In silico mixing experiment: four GLASS-OD samples initially classified as high-grade astrocytoma were incrementally spiked in with array data from non-tumor cortex and classified using MNP CNS v12.8. The x-axis represents the proportion of non-tumor data mixed in. The y-axis shows the median MNP class prediction probabilities for high-grade astrocytoma, oligosarcoma, and non-tumor cortex, with vertical bars indicating quantile distributions.

### No evidence for treatment-induced DNA methylation changes

As temozolomide and radiotherapy have been implicated in treatment-induced DNA alterations ^63,64^, we investigated whether these therapies leave detectable signatures in DNA methylation profiles. Due to the limited number of treated WHO grade 2 tumors (4 out of 80), differential methylation position analyses were restricted to the WHO grade 3 tumors. Comparing tumors treated with temozolomide (TMZ), any chemotherapy (TMZ, PCV, CCNU, BCNU, Cisplatin), or radiotherapy with untreated tumors identified 1, 2, and 0 significantly differentially methylated CpG sites, respectively (**Table S6**). These findings provide no evidence for apparent therapy-induced methylation patterns.

### Molecular mechanism of oligodendroglioma progression

We performed proteomics on 118 resections with matching methylation data available. To identify proteins associated with CGC^ψ^, linear regression was performed with CGC^ψ^ as continuous covariate. This resulted in 78/6.563 significant proteins (adj. p-value < 0.01; |LFC| > 0.5), enriched with proteins involved in (collagen-containing) extracellular matrix pathways (**Supplementary Fig. 9a**). Expression levels of MAG, MBP and PLP1, proteins implicated in myelination of oligodendrocytes, decreased when CGC^ψ^ increased, which could point towards dedifferentiation. Integration of the outcome with a similar proteomics analysis in the GLASS-NL astrocytoma dataset showed a correlated outcome (R=0.62), including up-regulation of collagen-containing extracellular matrix proteins and markers of tumor progression including PCNA and TMPO (**Supplementary Fig. 9b**). Both proteomics and the increased methylation of homeobox TFs suggest that tumor progression is associated with tumor cell de-differentiation.

DNA replication during cell division occurs with locus-specific timing. Using Repli-Seq, the relative timing of a locus being replicated during cell cycling (early/late replicon) can be measured. We examined whether the extent to which DNA methylation changed between WHO grades in oligodendrogliomas or with CGC^ψ^ was associated with early/late replicons, but did not observe a correlation (R=0.02 – 0.08; **Supplementary Fig. 9c**).

## Discussion

Making confident treatment decisions for patients with oligodendroglioma means balancing therapy intensity with quality of life, and to do that well, better ways to estimate how aggressive the tumor is are needed ^31^. To address this clinical need, we assembled a unique dataset from the GLASS consortium ^43^ including both primary and recurrent tumors, to investigate how DNA methylation patterns relate to tumor behavior. Out of this, we developed CGC^ψ^, a continuous scoring tool that is applicable to oligodendrogliomas and captures tumor malignancy in a more nuanced, objective way than traditional grading. We showed that CGC^ψ^ is robust and reproducible, and it could help define where along the spectrum of malignancy a tumor falls. Its prognostic value was better or as good as WHO grade. This application may be useful in the clinic by helping to determine when to escalate or de-escalate therapy, allowing treatment to be tailored to both prognosis and patient priorities. For example, it could be investigated whether there is a CGC^ψ^ threshold value up to which vorasidenib—assumed to be less effective in more aggressive tumors—provides benefit. This primary – recurrence study design also showed a notable DNA quality signal, at least partially due to time-dependent storage conditions. The effect was noticeable in samples stored in FFPE for 5 years or longer. This quality effect is probe and sequence-specific, typically affecting probes rich in G (guanine) or 5’ TA dinucleotides, which is in line with reduced probe binding due to cytosine deamination. The affected samples were characterized by lower overall intensity and differences in methylated/unmethylated channel intensity, even after normalization.

Beyond the quality effects, methylated levels change similarly — though with varying effect sizes — between primary – recurrent tumors, and between CNS WHO grades in oligodendrogliomas. However, the statistical power of the comparison between grades is considerably stronger than between the primary – recurrent resections, suggesting that this change does not take place in time linearly (as opposed to quality). This aligns with tumor evolution as a sequence of chance events but also implies that prognosis prediction of less aggressive tumors comes with more uncertainty.

Our results indicated that oligodendroglioma and astrocytoma progress along a shared prognostic epigenetic axis. However, astrocytomas and oligodendrogliomas, regardless of grade, remain distinguishable by their methylation profiles and do not evolve into an overarching malignant state. This axis constitutes at least a sequence context-specific decrease in methylation at sites with flanking sequences preferred by TET and increased methylation of polycomb transcription factor genes ^27^. We find the latter to be strongly correlated with epigenetic clocks, further in line with previous reports on the presence of a link between epigenetic aging and CNS WHO grade ^35^. Moreover, we demonstrate epigenetic aging at an accelerated pace in the light of CGC^ψ^, increased at tumor recurrence and linked to the Ki-67 positive cell fraction and density. This may be explained by a richer cell replicative history in more aggressive tumors. While the methylation profiles associated with tumor malignancy were a prominent source of variation in the data, we were not able to pinpoint a radio or chemotherapy-induced methylation profile, suggesting these treatments do not affect the methylome during course of the disease.

Our results corroborate a poor prognosis of oligosarcomas as they represent, like oligodendrogliomas classified as high grade astrocytomas, the extremes on the CGC^ψ^. The results further showed that the distinction between high grade astrocytoma and oligosarcoma is at least partially related to tumor purity, a factor the MNP CNS classifier is known to be sensitive for ^65^. The poor survival for tumors with a high CGC^ψ^, including but not limited to oligosarcomas, justifies prognostication of oligodendrogliomas based on DNA methylation ^25,65^. Both classifiers misclassified oligodendrogliomas as high-grade astrocytoma, specifically in aggressive recurrent cases with high tumor purity. For potential clinical applications, this should be taken into consideration.

Although we demonstrate that changes in DNA methylation were associated with patient prognosis, it remains to be determined whether this is causal or the effect of tumor progression and whether it takes place by passive or active DNA demethylation mechanism. The DNA demethylation showed CpG-specificity between the different oligodendroglioma datasets, as well as between astrocytomas. The demethylation is sequence context specific and correlated with TET enzyme flanking sequence preferences. This may be an indication for an active, TET-mediated, form of demethylation as underlying process to IDH-mutant glioma malignant transformation. This could further point towards reduced inhibition of TET enzymes by D-2HG ^66^, for instance by reduced levels of this onco-metabolite.

The *CDKN2A/B* homozygous deletion incidence was somewhat higher than the 6,83% in an external cohort (prospective French network; POLA) ^28^ and did not display a significant difference in survival from the last resection, although the mutated sample size was limited. In line with literature ^67^, hemizygous deletions typically have a large genomic span and therefore it is conceivable these losses’ selective advantage is not driven by *CDKN2A/B* disruption alone.

In summary, although several genetic and epigenetic molecular features are considered prognostic factors in oligodendrogliomas, grading these tumors remains difficult. We examined DNA methylation profiles of oligodendrogliomas and characterized their malignant phenotypes. The two most prominent changes, global demethylation and increased methylation of specific TFs, are shared with astrocytomas. Subsequently, we developed an astrocytoma derived continuous grading classifier that is applicable to and prognostic for oligodendrogliomas. This method is not based on nominal prognostic features such as infrequent DNA alterations, but on the basis of a continuous sequence context specific shift in the DNA methylation profile. The observed link between global demethylation and the preferential flanking sequences of TET demethylating enzymes is important, as it offers a potentially targetable explanation for the initiation and path of demethylation. Further research is needed to demonstrate its clinical relevance because of the retrospective nature of the study.

## Limitations of the study

This study is based on retrospectively collected data, which introduces heterogeneity in patient demographics and treatment regimens. While all patients had undergone two or more surgeries, some samples were excluded due to low purity or quality. For some patients either a primary or recurrent tumor sample was missing, limiting the ability to correct for patient-specific signals in those cases. The validation set is constrained by a limited number of samples with available survival data, variation in inclusion criteria across datasets, and differences in sample processing batches. Furthermore, it also included patients who underwent only a single surgical resection. Patients with multiple resections were preselected to have at least 6 months of overall survival and are generally expected to be fitter, which may introduce bias. Inclusion of both single- and multi-surgery patients could have resulted in a more heterogeneous overall survival distribution, reducing statistical power in survival analyses.

## Materials and Methods

### GLASS-OD discovery dataset

Patients histologically diagnosed with oligodendroglioma, IDH-mutant and 1p/19q-codeleted, who had undergone more than one surgical intervention with at least six months between the procedures, were eligible for inclusion. Tumor material and detailed clinical follow-up data were collected from institutes in Rotterdam, Amsterdam, Leiden, The Hague, and Utrecht (The Netherlands), as well as in Milan and Padova (Italy), and Durham (Duke University Medical Center [DUMC], USA). From DUMC, isolated DNA from fresh-frozen tissue was made available. From UMC Utrecht, processed DNA methylation arrays derived from FFPE material were available. Written informed consent was obtained from each subject. These studies were approved by the ethical board of the Erasmus MC (MEC-2020-0087, Rotterdam, The Netherlands), and conducted in accordance with institutional and national regulations.

### Validation dataset

A validation cohort was assembled from primary and/or recurrent oligodendrogliomas included from publicly available studies with distinct study designs ^11,30,44,45^. This cohort was complemented with three in-house oligodendroglioma samples resected only once. Samples were only included when they were processed using the Illumina BeadChip 850k V1 platform. As no sufficiently sized primary–recurrent EPIC 850k V1 dataset was available, we included oligodendroglioma samples regardless of the number of surgeries the patients had undergone. Based on CoNuMee CNV profiles, samples with insufficient estimated purity (<10%) or lacking 1p/19q-codeletion were excluded. WHO tumor grades were obtained from original pathology reports or corresponding manuscripts and were translated into CNS WHO grades using Arabic numerals instead of Roman numerals.

### Methylation array processing

For the majority of GLASS-OD samples, tissue was received as FFPE blocks or slides. The FFPE blocks were sectioned into 10 μm slices for DNA isolation, and 4 to 5 additional 10 μm sections were used for hematoxylin and eosin (H&E) staining. From these stainings, regions with the highest fraction of neoplastic cells were identified and marked on the sections by central neuropathologist JMK. These regions were then macrodissected and used for DNA isolation using the QIAamp DNA FFPE Tissue Kit. Both these samples and those received as isolated DNA were processed using Illumina’s Infinium MethylationEPIC v1.0 BeadChip (850k) at the internal facility of Erasmus MC.

### TCGA-LGG (1p/19q codel)

We obtained all TCGA-LGG *.idat files from https://portal.gdc.cancer.gov/projects/TCGA-LGG. Survival data and 1p/19q codeletion status were obtained from the literature ^46^. Only *.idat files from primary samples with matching entries labeled as 1p/19q codeleted were included. Additional survival data were obtained using TCGAbiolinks ^68^. Survival data from TCGAbiolinks were converted from months to days by multiplying by 30.43686. Due to incomplete survival data in both sources, survival data (in days) were intersected. In cases where survival data were available from both sources, the TCGAbiolinks data were used, as they were more up to date (**Table S7**). In total, 150 *.idat samples with survival data were available, of which 21 had a recorded survival event.

### MNP CNS Classifier

Data were classified via the MNP portal using the Predict-Brain classifier ^36^. The CoNuMee package, as incorporated in MNP classifier v12.8, was used to determine copy number variations (CNVs). Quality control (QC) metrics were obtained from classifier version v11b4. For 10 out of 121 patients in the discovery dataset, resections were misclassified as astrocytoma due to the absence of a detectable 1p/19q codeletion. These cases were excluded from all subsequent analyses. The final GLASS-OD methylation cohort consists of 211 resections from 111 patients.

### GLASS-NL dataset

Astrocytoma, IDH-mutant samples from the GLASS-NL study were obtained ^35^. Samples with a fraction of detPfailed probes greater than 2.5% were excluded, resulting in 219 samples retained for analysis. These were used for principal component analysis (PCA). Principal component 3 (PC3) correlated with tumor purity and segregated samples with a flat CNV profile. Of the 219 samples, those with a PC3 value below 300 were retained (n = 203).

### DNA-methylation processing and analysis

To evaluate sample and probe quality, all *.idat files were loaded using minfi 3.18 ^69^ in R and pre-processed with offset=0, dyeCorr=T and dyeMethod=“single”. Dye correction was set to single to ensure normalization outcome is independent of the poorest quality sample of the run(s) and comparable between runs. While minfi loaded all n=865.859 probes, only the n=760.405 probes not annotated as MASK_general = TRUE were exported to M-values. Detection-P (det-P) fractions were estimated using: detectionP(type = “m+u”). Probes with an insignificant p-value (pval > 0.01) were marked as failed and the per-probe and per-sample fraction of failed probes was estimated. Low-performing samples and probes were identified by a combination of per-sample det-P failed probes and principal component analysis used only for QC analysis, upon the 200.000 most variable (stats::mad) M-values. By visualizing the per sample percentage det-P value with the correlated principal component, PC1, we exclude samples with a det-P fraction of >= 0.025 probes or a PC1 >= 875. After tumor purity and quality control, DNA methylation data was available for 229 resections from 121 patients. Copy-number plots were examined for evidence of 1p/19q codeletion. No codeletion was detected in 18 samples from 10 patients, and these cases were excluded from further analysis.

Of these 211 samples and n=685.271 probes passing quality control, resulting m-values (minfi::ratioConvert(…, what = “M”)) were exported to a cache file. All post-qc analyses were only conducted on these n=685.271 probes that had not failed detection-P, PCA, targeted specifically CpGs, of the 760.405 probes not annotated as MASK_general = TRUE, except for external tools which required all probes for input (epigenetic clocks and MNP classifier). Beta-values were estimated (minfi::ratioConvert(…, what = “beta”)) for external epigenetic clock packages and estimation of per-sample mean/median methylation levels. For sample-to-patient fingerprinting analysis, beta-values of SNP covering probes were extracted (minfi::getSnpBeta(RGSet)) and scaled (scale(, center=F)) before calculating the sample-to-sample Euclidean distance. A post-qc principal component analysis (PCA) was performed on the M-values of GLASS-OD and GLASS-NL datasets, separately. The heatmap including all 1p/19q surgical interventions was clustered on these principal components PC2 - PC20. PC1 was visualized but excluded from this clustering because of its strong associations with quality. UMAP was performed on the top 7,500 most variable CpG probes (mean absolute deviation), not located on chromosomes X, Y and M, using the uwot library. Samples from the validation set were mapped onto the GLASS-OD PCA using the *predict()* function in R. For gene-level interpretation of DNA methylation, the mean t-statistic for all DNA methylation probes per gene were compared with the distribution of the mean t-statistics for the remaining genes. Linear regression analysis was performed using limma to find per-bin copy-number variations associated with malignant progression. In the CNV models, tumor purity was incorporated as continuous covariate in the respective regression models as well as per-patient correction. GLASS-OD samples were clustered using ComplexHeatmap ^70^. Tumor purity estimates were plotted along their CoNuMee/CNVP estimated profile. Raw *.idat files and processed M-values are available at GEO under accession GSE297733.

### Tumor purity

For oligodendrogliomas tumor purity was estimated using the CNV foldChange (Connumee CNVP v5.2 segment calls) for the 1p/19q codeletion. The respective foldChanges for all genomic bins at chromosomal arms 1p and 19q were used for a 1p/19q codeletion specific principal component analysis, used to select features for purity estimation. The first component was associated with the intensities of the codeletions from both chromosomal arms. From these bins, only the 1.013 bins with PC1 loadings between -0.021 and - 0.0345 were selected (**Table S8**). Then from these bins strongly contributing to PC1, the median foldChange *f* was calculated. Tumor purity *p* was defined as: *p* = –2 * (2^*f*^ – 1). Array samples with a purity below 0.1 (10%), of which none was classified as oligodendroglioma by MNP, were excluded from further analysis. In the validation cohort, 5 samples were excluded due to low purity (**Supplementary Fig. 1a**). For astrocytomas, PC3 related to tumor purity as it segregated samples with a flat CNV profile. Of the 219 samples, those with a PC3 value smaller than 300 were kept (n=203).

### Epigenetic clocks

Prior to running epigenetic clock algorithms, samples were first normalized (minfi::preprocessNoob(RGSet, offset = 0, dyeCorr = T, dyeMethod=“single”)), exported to beta-values and inserted in epigenetic clock algorithms using the dnaMethyAge metapackage ^61^, including the following clocks: HannumG2013 (human whole blood), LevineM2018 (human whole blood), ZhangQ2019 (human blood and saliva), ShirebyG2020 (human cortex), ZhangY2017 (fitted to time to death; human blood of patients with 38 diseases, including cancer), LuA2019 (human whole blood), HorvathS2018 & PCHorvathS2018 (multiple human cell types; non cancer), McEwenL2019 (buccal epithelial cells), CBL_specific, PCHorvathS2013 (multiple human cell types; non cancer), PCHannumG2013 (human whole blood), PCPhenoAge, CBL_common, Cortex_common, LuA2023p1 & LuA2023p2 & LuA2023p3 (multiple tissue types across multiple mammalian species), YangZ2016 (whole blood). EpiTOC2 ^60^ and RepliTali ^71^ were executed using their original package.

### CGC^ψ^

CGC^ψ^ was generated as LASSO model (glmnet library) and was trained on the GLASS-NL dataset with CGC, formally log(A_IDH_LH / A_IDH_HG) ^25^, as response variable, with alpha = 1 and lambda = 0.1041977. The input for CGC^ψ^ prediction was the M-value matrix of the 685,271 probes (offset = 0, dyeCorr = TRUE, dyeMethod=“single”) passing quality control, of the 203 GLASS-NL samples passing quality control. For the TCGA-LGG dataset, a derivative model was trained using only the intersection of 685,271 850k probes and 450k array probes. This model was then applied to the TCGA-LGG 1p/19q samples only. A small R package was written to apply the linear predictors: https://github.com/ErasmusMC-Neuro-Oncology/Continuous_Grading_Classifier/

### Mixing idat files: idat-tools

For mixing oligodendroglioma *.idat files classified as high grade astrocytoma by the MNP CNS classifier with incremental fractions of non-tumor sample, we developed a free open-source software application in python3, idat-tools: https://github.com/yhoogstrate/idat-tools/. It can read an *.idat file into memory, change its contents, and write the memory object back to a new *.idat file. By mixing the *.idat file with a second *.idat file, given a fraction (0.0 – 1.0), all data, including idat columns std_dev, n_probes and intensity, are proportionally mixed, rounded and exported. For *.idat files not having an identical number of probes, the intersection of probes is taken. In this study, samples were mixed with fractions {0.1, 0.2, … 0.8, 0.9} (**Table S5**).

### Differential methylated probe analysis

DMP analyses were performed by (typically multi-variate) linear modeling of the M-values using limma ^72^. M-values were chosen instead of beta-values due to their symmetrical distribution centered around zero and their unbounded range, both of which make them more suitable for linear modeling. Because earlier DNA methylation profiling of oligodendrogliomas reported the presence of patient-specific effects ^73^, we aimed to factor these out by incorporating patient identifiers in multivariate models. For each experimental design, the included single-patient/patient-unique samples were group into a decoy “remainder” patient. Multivariate models were generated using model.matrix (e.g. model.matrix(∼factor(patient) + factor(primary.recurrence), data=…)), then fitted using lmFit and tested using limma::eBayes(…, trend=T). The appropriate coefficients were exported with limma::topTable (e.g. limma::topTable(…, n=nrow(…), coef=“factor(primary.recurrence)recurrence”, sort.by = “none”, adjust.method=“fdr”)). For visualizations, the subsequent t-statistics were used as they are signed and continuous unlike p-values, and corrected for standard error, unlike the log2foldChange. CpG probes were considered significantly different with an FDR adjusted p-value < 0.01 and a |log2FC| > 0.5. For multivariate models, incorporated continuous factors such as CGC^ψ^ were typically scaled to a standard deviation of 1.

### Polycomb gene annotations

Genes attributed to Polycomb complexes were taken from literature ^74^.

### Gene enrichment

The GencodeCompV12_NAME column from Illumina’s infinium-methylationepic-v-1-0-b5-manifest-file.csv manifest was used for probe-to-gene annotation. For probes annotated to more than one gene, the annotation was split and the table was expanded. Genes starting with “RP” followed by numbers were excluded. For each gene, the median t-statistics (between WHO grades) of all probes belonging to the gene was computed and compared with the median t-statistics of other genes. Genes with HUGO symbols starting with ‘OR’ followed by a number (regex: “^OR[0-9]”) were considered olfactory receptor family genes.

### RepliSeq

Processed replication timing experiment results were downloaded from UCSC’s goldenpath wgEncodeUwRepliSeq track ^75^ (bigWigs files, hg19). These were converted to hg38 using CrossMap ^76^ and exported to bedgraph. CpGs were annotated with their respective replication-seq timing value by intersection with the lifted-over bigWig bins using their genomic coordinates and the HTSeq library ^77^ in python3. Excessive outliers in RepliSeq score were excluded (keep: BjWaveSignalRep2 < 79; Bg02esWaveSignalRep1 < 90; BjWaveSignalRep1 < 95; BjWaveSignalRep1 > 10; NhekWaveSignalRep1 < 80). Spearman correlation was estimated between the per-CpG change in methylation between WHO grades with the respective RepliSeq values.

### Sequence Contexts

Probes were mapped to their sequence context deduced from “infinium-methylationepic-v-1-0-b5-manifest-file.csv”. Probe sequences “AlleleA_ProbeSeq” were compared with the genomic alignment sequence “Forward_Sequence” to determine whether the probe’s orientation is forward. The Forward_Sequence was used to extract the sequence context and when probes are in reversed orientation, the reverse complement of the Forward_Sequence was used. TET and DNMT enzyme flanking sequence preferences were combined from literature ^51–58^. Both t-statistics and enzyme affinities were integrated from their 256 stranded sequence context into 136 unstranded contexts by taking their mean.

### Ki-67 staining and quantification

Tissue slices were used for Ki-67 staining with respective control tissue (tonsil) appended on the slide. Immunohistochemistry was performed with an automated, validated and accredited staining system (Ventana Benchmark ULTRA, Ventana Medical Systems) using the ultraview universal DAB Detection Kit. In brief, following deparaffinization and heat-induced antigen retrieval the tissue samples were incubated according to their optimized time with the antibody of interest (**Table S9**). Incubation was followed by detection with the secondary antibody included in the untraview universal kit (multimere), followed by haematoxylin II counter stain for 20 min and then a blue coloring reagent for 8 min according to the manufacturer’s instructions (Ventana Medical Systems Inc., Arizona, USA). The scanned stainings were imported in QuPath 0.5.0 (*.ndpi files).

Samples were imported as Brightfield H-DAB. Regions containing tissue were manually selected and artifacts (gaps and folds) were manually excluded. Within these regions cells were detected using StarDist2D (dsb2018_heavy_augment; dsb2018_paper; he_heavy_augment; normalizePercentiles: 1, 99); threshold: 0.40; pixelSize: 0.2276; includeProbability: true; cellExpansion: 5). A total of n=20 samples were used for training cell type classification, in which a fraction of cells were selected and annotated as either Ki67+, Ki67- or “other” in case of erythrocytes, air bubbles or other artifacts. A classifier was trained using TrainObjectClassifier on all training cells (Classifier: ANN_MLP; all measurements; all 3 classes). This classifier was exported as *DABnn6* (https://github.com/yhoogstrate/glassod/tree/main/QuPath/classifiers/object_classifiers). The classifier was applied to all regions runObjectClassifier(“DABnn6”);. Data were exported as *measurements*. Snapshots were exported from QuPath to ImageJ in which the scale bars were added.

### Proteomics

From 140 unique resections, two 10µm FFPE tissue sections were used for proteomics analysis. The system used was a Bruker timsTOF attached to an EvoSep system using dia-PASEF ^78,79^. Of these samples, 118 had matching DNA methylation data. Per-peptide intensities were transformed into 8.070 protein-wise intensities using prolfqua ^80^. Control entries and entries lacking HUGO gene annotations were excluded. Proteins with more than 50% N/A values were excluded. Intensities were log2 transformed and per-sample robust scaled using median and IQR. In total, 6.540 proteins were available for downstream analysis. Limma ^72^ was used for statistical comparison with CGC^ψ^ as a continuous factor. After CGC^ψ^ was scaled using the scale function, protein expression was considered significant with an FDR-adjusted p-value < 0.01 and |log2FC|>0.5.

### R statistical computing

R v4.4.2 was used for data processing, including the limma, uwot, tidyverse, patchwork, survival, survminer, ggrepel, ggbeeswarm, factoextra, pROC, glmnet and ggpubr libraries. The code is freely accessible at the following repository: (https://github.com/yhoogstrate/glass-od).

## Data availability statement

Raw *.idat files and processed M-values will become available at GEO under accession GSE297733. Analysis R code is freely accessible at the following repository: https://github.com/yhoogstrate/glass-od. A small R package was written to apply the linear predictors: https://github.com/ErasmusMC-Neuro-Oncology/Continuous_Grading_Classifier/

## Acknowledgements

We kindly acknowledge Dr. Abigail Suwala for making details and/or data available for completing the validation set. We also kindly acknowledge the Functional Genomics Center Zurich (FGCZ) of University of Zurich and ETH Zurich and Erasmus MC Pathology Research and Trial Service (PARTS) for their services and using their facilities.

## Author contributions

Conceptualization: Y.H.; S.A.G.; L.H.; M.P.; A.D.; M.M.J.W.; R.L.; T.W.; M.C.M.K.; Y.K.; B.Y.; P.W.; P.J.F.; Formal analysis and investigation: Y.H.; S.A.G.; L.H.; R.H.; I.H.; M.P.; M.d.W.; A.D.; A.B.; F.S.V.; Resources: all (co-)authors; Writing (initial): Y.H.; S.A.G.; P.W.; P.J.F.; Writing – review & editing: all (co-)authors; Supervision and funding: Y.H.; P.W.; P.J.F.

---

**Supplementary Fig. 1.**
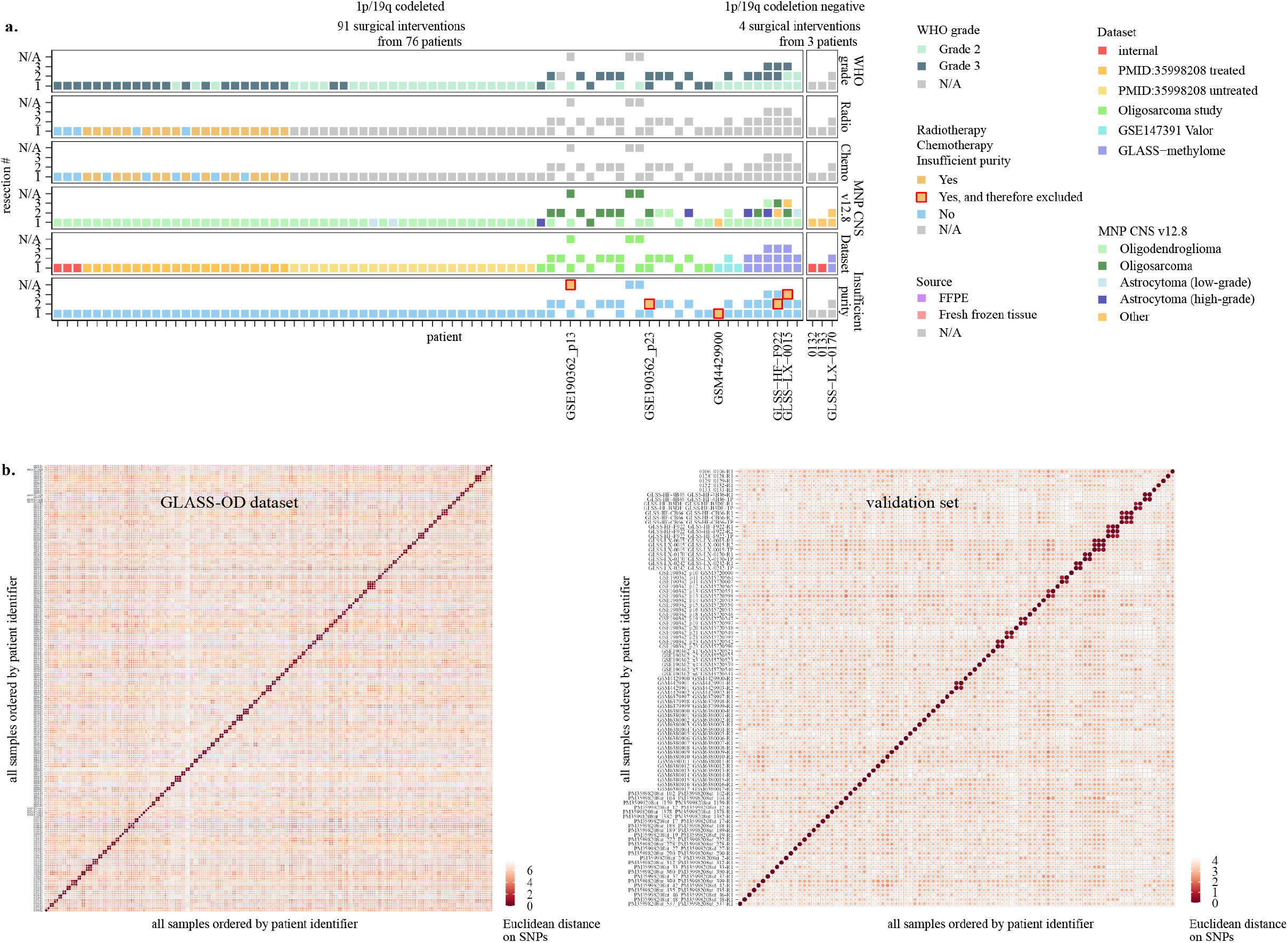
Cohort overview and quality control analyses. **(a)** Patient overview for the validation DNA methylation dataset, analogous to Fig. 1a. Samples excluded due to low purity or absence of 1p/19q codeletions are labeled. **(b)** SNP-probe-based fingerprinting. Both axes represent all samples per dataset, ordered by patient identifier and then surgery number. Each dot indicates the Euclidean distance between two samples based on scaled beta-values from SNP probes. Light colors indicate large distances between two samples (low similarity), while dark colors indicate small distances between two samples (high similarity). Samples that belong to the same individual are represented by multiple dark dots.

**Supplementary Fig. 2.**
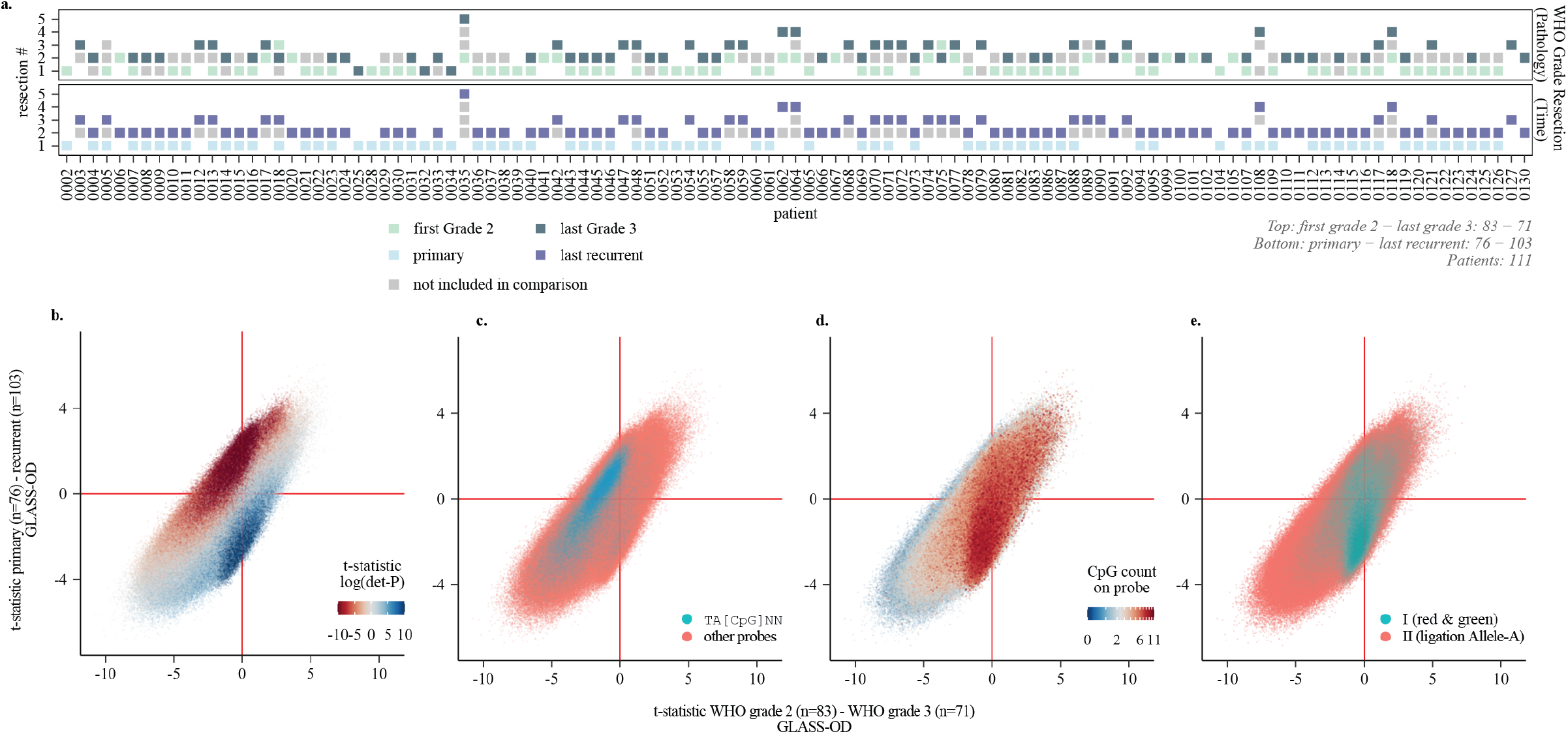
Notable effect of data quality on DMP outcomes. **(a)** Distribution of samples across groups used in the two DMP comparisons. Top: comparison between the first CNS WHO grade 2 samples (light green) and the last WHO grade 3 samples (dark green). Bottom: comparison between primary surgical interventions (light blue) and the last surgical interventions (dark blue). **(b)** Same integrated DMP plot as in Fig. 2c, with probes colored by the t-statistic from an additional DMP model relating CpG methylation levels to the log-transformed percentage of detection p-value failed probes per sample. **(c)** Same integrated DMP plot as in Fig. 2c, with probes colored by having a TA[CpG]NN sequence context. **(d)** Same integrated DMP plot as in Fig. 2c, with probes colored by the total number of CpG sites within the probe sequence. **(e)** Same integrated DMP plot as in Fig. 2c, with probes colored by the probe chemistry type.

**Supplementary Fig. 3.**
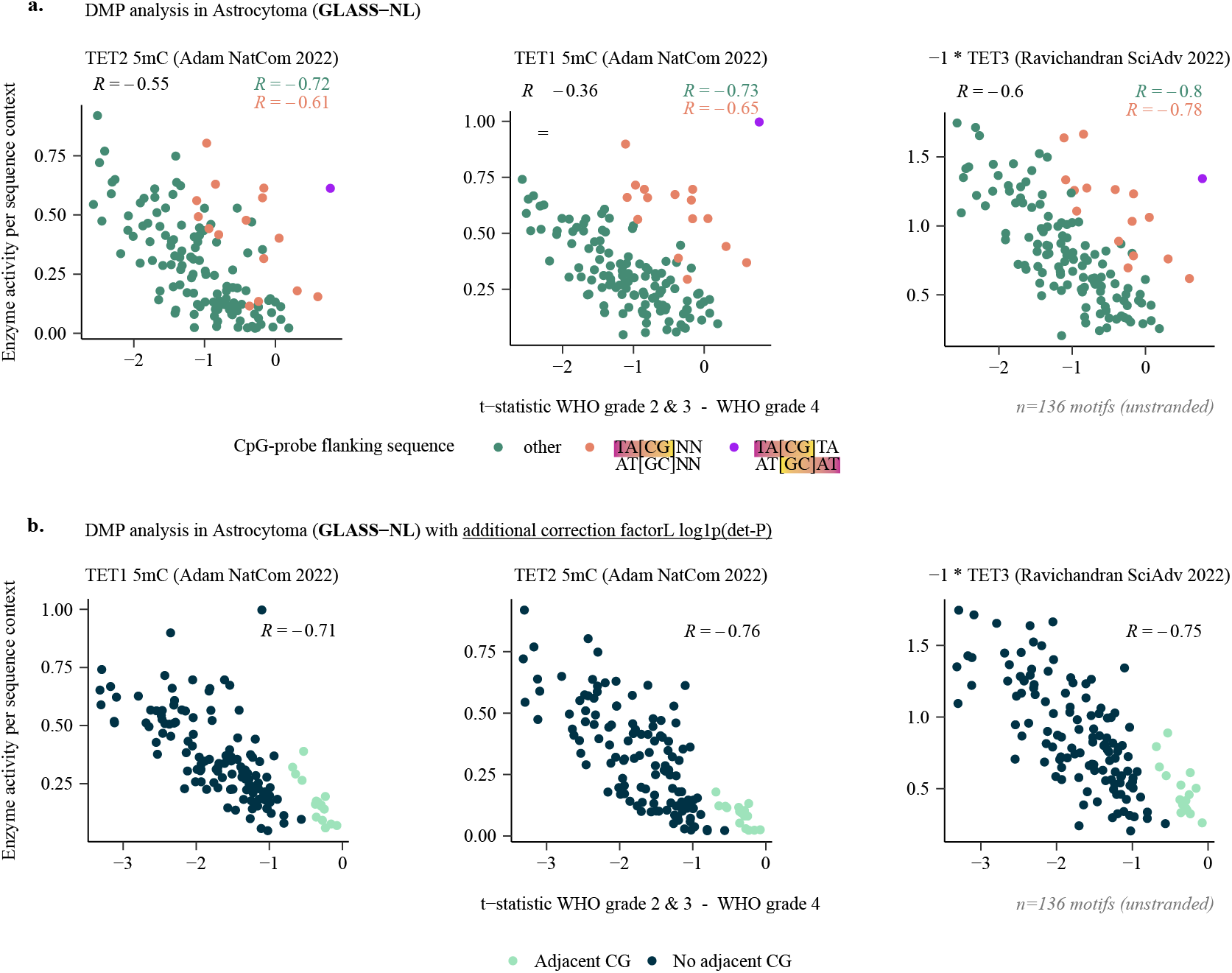
Changes in methylation are sequence context specific in astrocytoma (GLASS-NL). **(a)** Scatterplots showing the mean t-statistics per sequence context (x-axis), comparing methylation differences between WHO grade 2 & 3 versus grade 4 astrocytomas, plotted against TET flanking sequence preference (y-axis). Sequence contexts are color-coded by matching the quality associated TA[CpG] subsequence once, twice (palindromic) or not. Spearman correlation coefficients (ρ) are shown in black for all data points, in orange for contexts with one TA[CpG] subsequence, and in green for those not matching. **(b)** Same as (a) but using a model correcting with log1p(fraction det-P failed probes).

**Supplementary Fig. 4.**
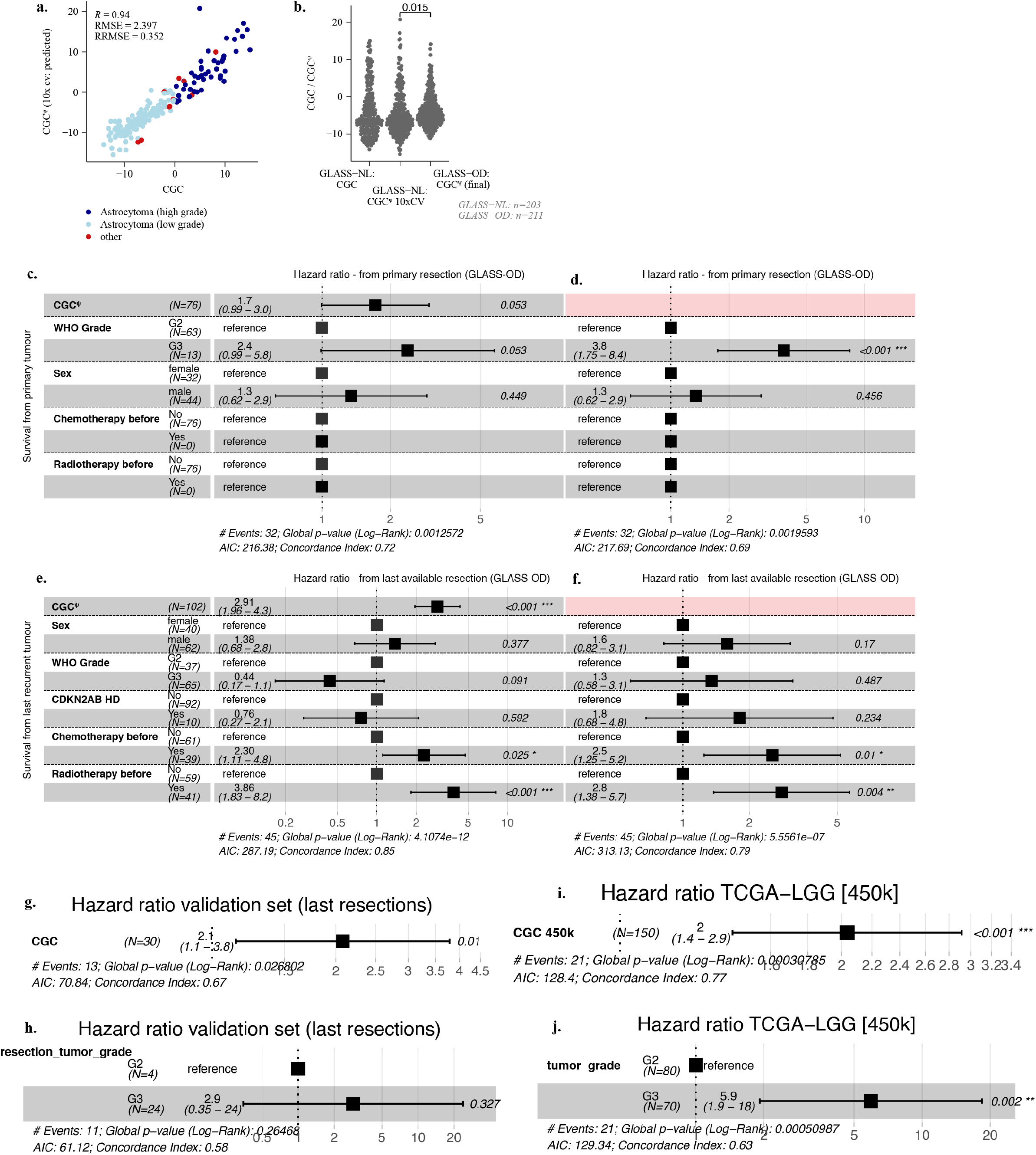
CGC^ψ^ model development and multivariate survival analysis. **(a)** Scatterplot showing the expected CGC values (actual CGC) versus the observed CGC^ψ^ estimates obtained through 10-fold cross-validation in all GLASS-NL samples. **(b)** Density scatterplots showing: (left) actual CGC scores in the GLASS-NL dataset, (middle) predicted CGC^ψ^ scores from 10-fold cross-validation in GLASS-NL, and (right) CGC^ψ^ scores from the final model applied to oligodendrogliomas in the GLASS-OD dataset. The difference in CGC^ψ^ between samples from the GLASS-NL and GLASS-OD datasets was test using the Wilcoxon rank-sum test, shown at the top. **(c)** Forest plot of multivariate survival analysis for overall survival of primary tumors, including both CGC^ψ^ and CNS WHO grade as covariates. **(d)** Forest plot of multivariate survival analysis for overall survival of primary tumors, including CNS WHO grade but excluding CGC^ψ^. **(e)** Forest plot of multivariate survival analysis for post-recurrence survival of the last available tumors, including both CGC^ψ^ and CNS WHO grade. **(f)** Forest plot of multivariate survival analysis for post-recurrence survival of the last available tumors, including CNS WHO grade but excluding CGC^ψ^. **(g)** Forest plot of univariate CGC^ψ^ in the last available resections of the validation set. **(h)** Forest plot of univariate WHO grade in the last available resections of the validation set. **(i)** Forest plot of univariate validation of CGC^ψ^/450k in the primary resections of oligodendroglioma samples from the TCGA-LGG dataset (450k arrays). **(j)** Forest plot of univariate validation of WHO grade in the primary resections of oligodendroglioma samples from the TCGA-LGG dataset (450k arrays).

**Supplementary Fig. 5.**
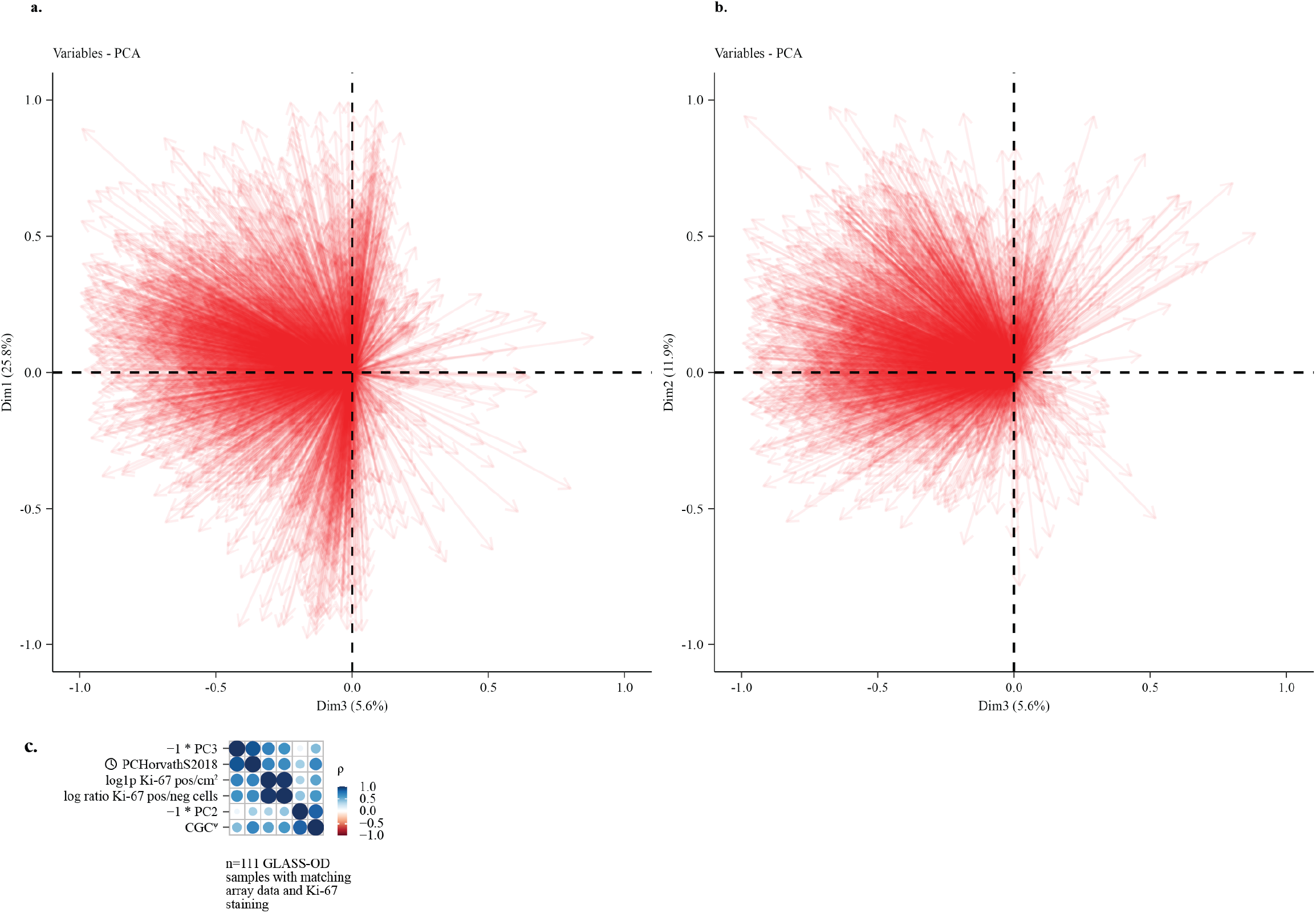
CpG probes annotated to belong to polycomb transcription factors contribute to PC3. **(a)** Variables factor map for principal components 1 and 3, marking only probes mapped to polycomb transcription factors. **(b)** Variables factor map for principal components 2 and 3, marking only probes mapped to polycomb transcription factors. **(c)** Spearman correlation (ρ) based clustering of Ki-67 positive cell ratio and density with other sample parameters, for the n=111 samples with matching Ki-67 staining and DNA methylation arrays.

**Supplementary Fig. 6.**
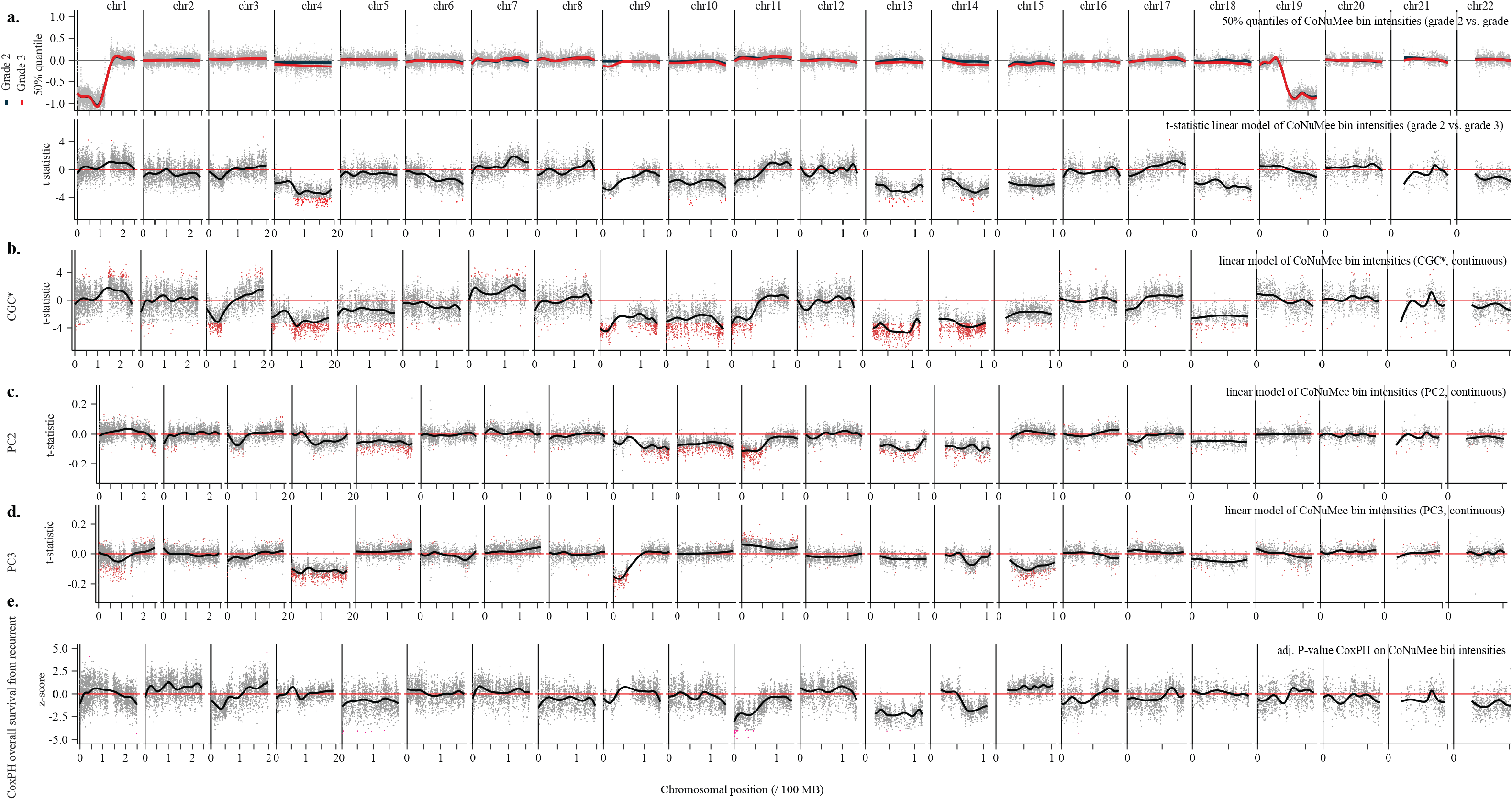
Recurrent tumors are characterized by chromosomal losses. **(a)** Differences in intensity per CNV bin compared between WHO grades in oligodendroglioma. **(b)** Differences in CNV bin intensity associated with CGC^ψ^. T-statistics and FDR-adjusted p-values are derived from multivariate linear models fitted to CNV bin intensity, including both tumor purity and CGC ^ψ^ (scaled to a standard deviation of 1) as covariates. Regions with a significant association (p<0.01) are marked in red. **(c, d)** Same as (b), but using including tumor purity, PC2 and PC3. Regions with a significant association (p<0.01) are marked in red. c00000ff Same as (b), but using a CoxPH model on overall survival at tumor recurrence. Regions with a trend in association (p>=0.01 & p<0.05) are marked in pink.

**Supplementary Fig. 7.**
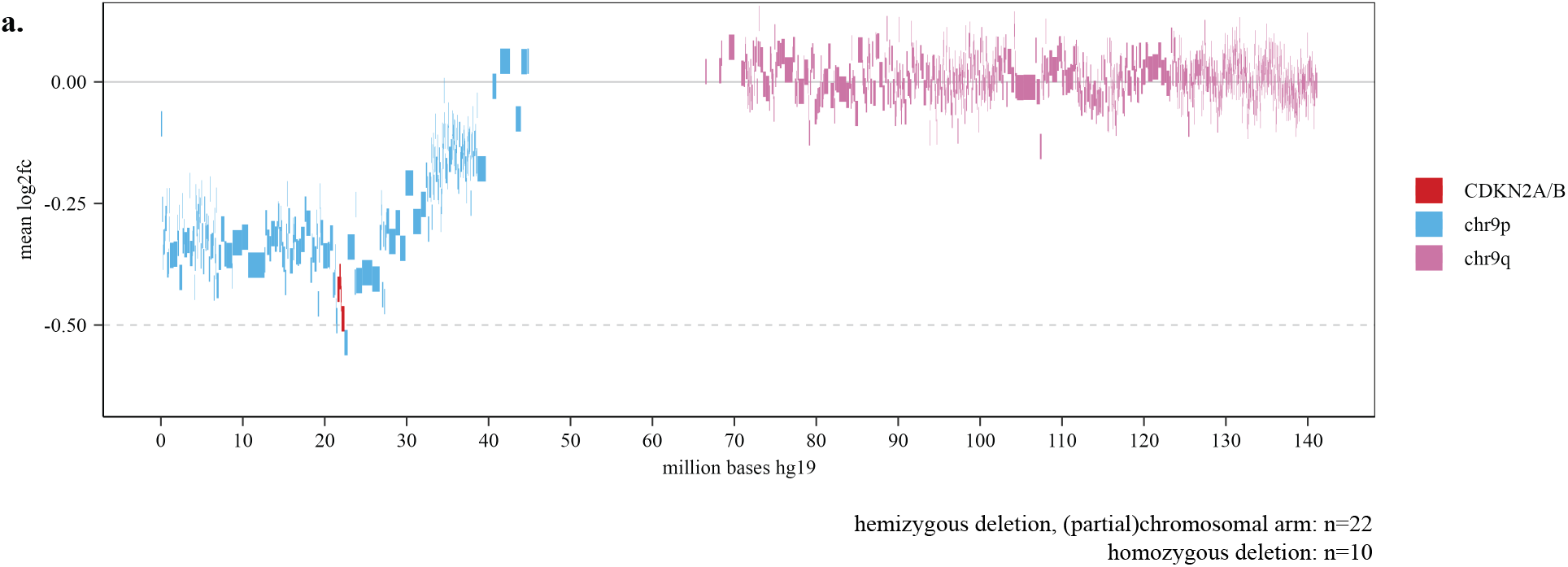
The *CDKN2A/B* locus is frequently deleted as a result of partial loss of the chr9p arm. **(a)** Detailed copy-number view of chr9, showing the average CoNuMee score of samples with a chr9p arm, partial chr9p arm or a homozygous deletion of *CDKN2A/B* (n=32; **Table S4**).

**Supplementary Fig. 8.**
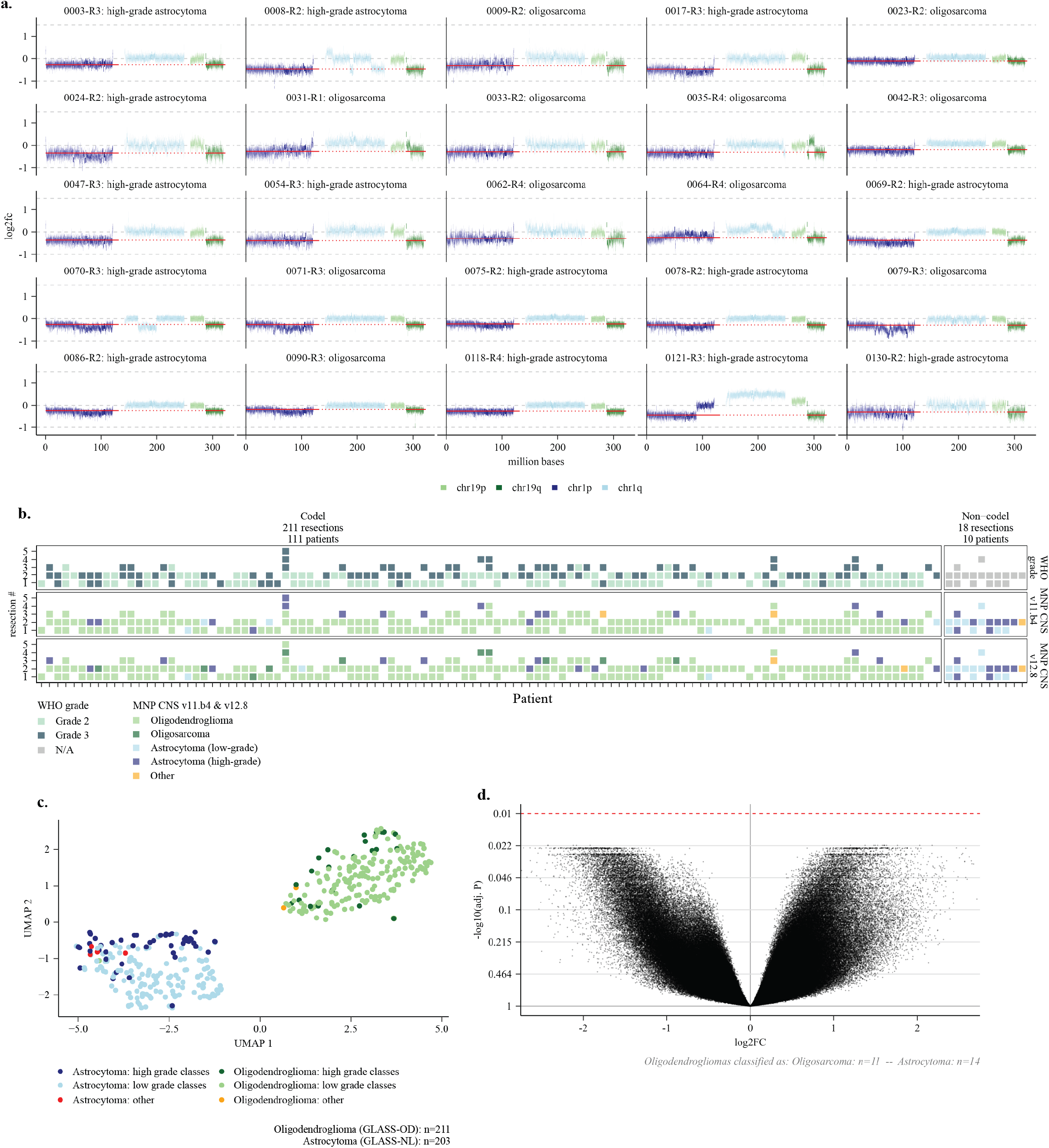
High grade oligodendrogliomas and astrocytomas remain epigenetically distinct tumor types. **(a)** CNV profiles of chromosomes 1 and 19 for GLASS-OD samples classified as either high-grade astrocytoma or oligosarcoma. Chromosome arms are color-coded to highlight the presence of the 1p/19q co-deletion. The x-axis shows genomic distance along chr1 or chr19. The y-axis represents the foldchange as provided by CoNuMee. Red lines indicate tumor purity estimates (based on bin intensities of 1p and 19q). **(b)** Same patient overview as in Fig. 1a, extended with classifications from MNP CNS classifier v11.b4. **(c)** Uniform Manifold Approximation and Projection (UMAP) of samples from the GLASS-OD (n=211) and GLASS-NL (n=203) datasets, colored by MNP CNS classifier grade groups: low-grade (light colors; low-grade astrocytoma or oligodendroglioma) and high-grade (dark colors; high-grade astrocytoma or oligosarcoma). **(d)** Volcano plot showing differential methylation between oligodendrogliomas classified as high-grade astrocytoma (n=14) or oligosarcoma (n=11) by the MNP CNS classifier. Each dot represents a CpG probe. The x-axis represents the log_2_ fold change, and the y-axis shows the –log_10_ FDR-adjusted p-value. The red dashed line represents the adjusted p-value cut-off of 0.01.

**Supplementary Fig. 9.**
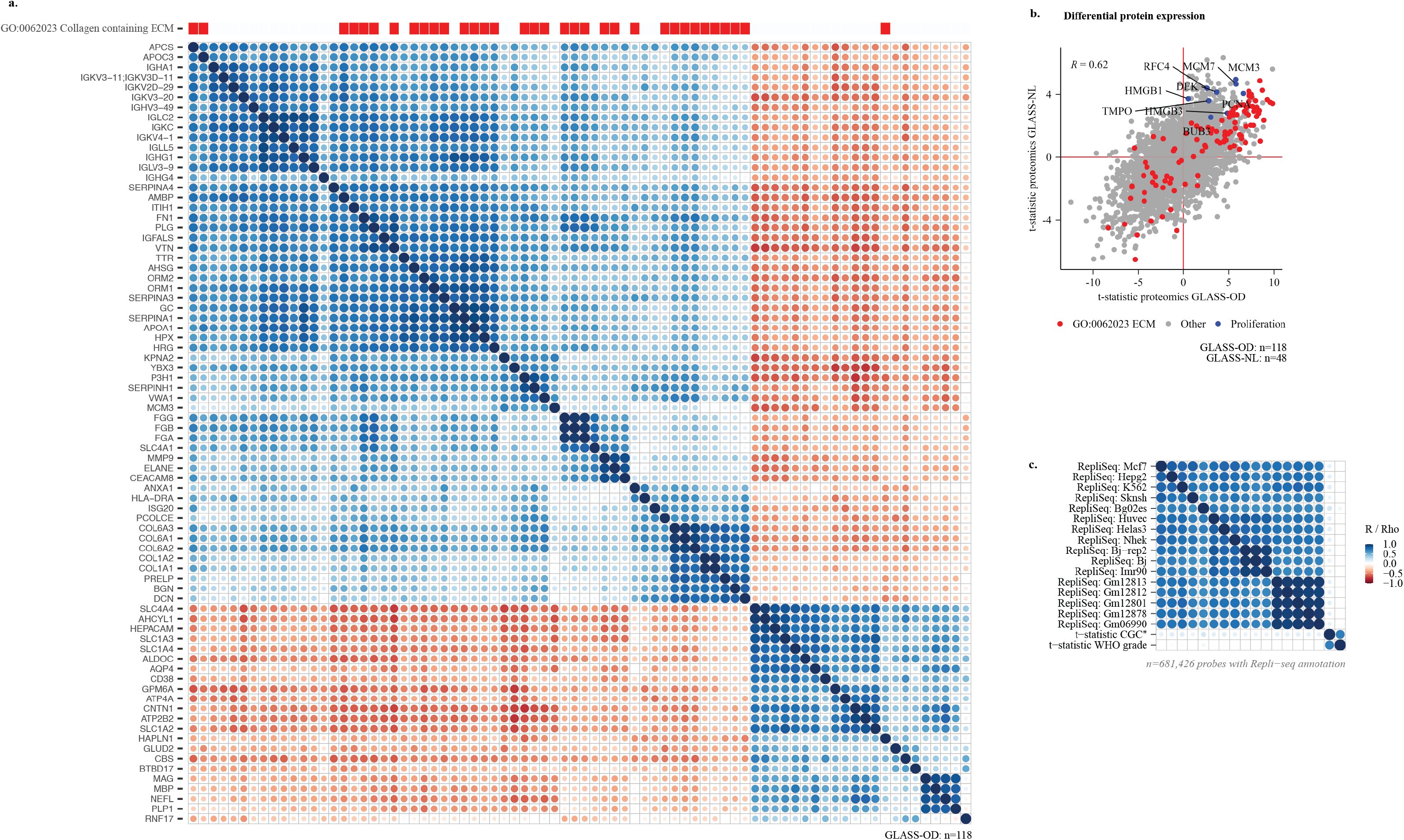
Collagen ECM-related protein expression and replication timing in relation to CGC^ψ^. **(a)** Pearson correlation-based clustering of proteins significantly associated with CGC^ψ^. Proteins corresponding to genes in the collagen-containing extracellular matrix (ECM) pathway are highlighted at the top. **(b)** Intersection of differential protein expression results between oligodendrogliomas from GLASS-OD (fitted to CGC^ψ^, x-axis) and astrocytomas from GLASS-NL (fitted to CGC, y-axis). Proteins associated with the collagen-containing ECM are marked in red, commonly used proliferation markers are marked in blue. Pearson correlation coefficient (R) is indicated. **(c)** Correlation between RepliSeq-based replication timing (per genomic bin) and median t-statistics comparing WHO grades in oligodendrogliomas (per same genomic bin). Spearman’s rank correlation coefficient (ρ) is shown.

